# Behavioral and phenotypic constraint belie deep genomic divergence and seasonal adaptation in a widespread desert lizard

**DOI:** 10.1101/2025.09.29.679313

**Authors:** Raúl Araya-Donoso, Elizabeth Dávalos-Dehullu, Zachary W. Lukasik-Drescher, Douglas G. Moore, Benjamin T. Wilder, Andrés Lira-Noriega, Adrián Munguía-Vega, Kenro Kusumi, Greer A. Dolby

## Abstract

Cryptic species offer opportunities to reveal the mechanisms that constrain phenotypic divergence during speciation. We integrated whole-genome sequencing, morphological, micro- and macro-climatic, and behavioral data to investigate divergence across a well-documented genetic break in the desert-adapted side-blotched lizard, *Uta stansburiana,* on the Baja California peninsula. Despite deep genomic differentiation, clades show remarkable similarity in morphology, habitat use, and thermal biology. Nearly all genetic differentiation (87%) is explained by isolation by distance and seasonal variation in precipitation, with almost no effect of temperature. Behavioral thermoregulation and changes in activity time accommodate strong macro- and micro-climatic differences, buffering against selection that would otherwise drive morphological and physiological divergence. In contrast, genomic signatures of selection and divergence in genes associated with the nervous system, sensory perception, and biomolecule metabolism indicate adaptation to differences in rainfall seasonality. The results show behavioral flexibility can constrain phenotypic divergence, yielding cryptic species-level genetic divergence despite strong eco-climatic disparities and selection pressures. More broadly, this study shows how rigorous statistical integration of multiple data types can disentangle competing eco-climatic drivers that can decouple phenotype from genotype during speciation.

**Significance:** Understanding why deep genetic divergence occurs without phenotypic differentiation is a longstanding challenge in evolutionary biology. By statistically integrating genomic, morphological, climatic, and behavioral data, we test the mechanisms controlling differentiation within a natural lizard system in a geo-climatically diverse setting. Results show that isolation by distance and adaptation to precipitation seasonality drive nearly all genomic differentiation. Behavioral adjustment to strong thermal variation buffers against selection pressure otherwise expected to cause divergence in morphology, thermal biology, and habitat use. This work demonstrates how rigorous integrative analyses can tease apart ecological and neutral factors controlling genomic divergence, providing rare insight into causal mechanisms driving speciation while constraining phenotypic divergence.

## Background

A pivotal challenge in evolutionary biology is understanding the mechanisms that generate differential patterns of species richness, genetic diversity, and phenotypic diversity (1).

Some groups of organisms, like those in adaptive radiations, have rapid accumulation of genetic and phenotypic differentiation that results in high rates of species formation (2). In contrast, others accumulate high genetic differentiation without phenotypic disparity (e.g. cryptic species and non-adaptive radiations). The study of cryptic species provides an opportunity to explore the intrinsic and extrinsic factors that constrain phenotypic differentiation in the face of divergent selection pressures and speciation (3, 4).

During speciation, the accumulation of genetic and phenotypic divergence are not necessarily coupled (5). Young age of lineage split, convergent evolution, developmental constraints, or low evolutionary potential and low divergent selection pressures can explain the lack of phenotypic differentiation within cryptic species (4). The strength of selection can be reduced if there is strong genetic drift at play (e.g. low *N_e_*) (6). Alternatively, organisms can accommodate selection pressures through phenotypic, behavioral, and epigenetic plasticity instead of adaptation *per se*, defined as underlying changes in allele frequencies (7). For example, ectotherms can meet their optimal temperatures through behavioral adjustments, reducing the *effective* selection pressure on thermal physiology (8). Finally, strong purifying selection can also limit phenotypic differentiation if aspects of the phenotype are constrained by development or ecology (4).

The integration of Whole Genome Sequencing (WGS) data with morphological, habitat use, and behavioral data can help disentangle the complex mechanisms that decouple genetic and phenotypic divergence (9). For instance, it enables the assessment of neutral and adaptive processes shaping genomic evolution (10), and by identifying putatively adaptive genomic mutations we can infer which aspects of the phenotype are responding to selection pressures and which may not be. Collection of microhabitat-use and behavioral data can reveal how an organism responds to different macro-climates. The combination of this field-based natural history information with genomic data is particularly useful when the phenotypic traits under divergence cannot be observed or measured.

In this study, we assessed genomic, phenotypic, and eco-behavioral differentiation in the side-blotched lizard, *Uta stansburiana*. The genus *Uta* is a good model because although it diverged from its sister genera *Urosaurus* and *Sceloporus* ∼40 Mya (11), only one species (*U. stansburiana*) has been described for continental areas, and it is widely distributed across southwestern North America (∼2,700,000 km^2^). It has six island-endemic species (12), which suggests that there may be factors constraining its morphological differentiation on the continent. In contrast, *Sceloporus* and *Urosaurus* have 118 and 8 species, respectively (12). *Uta stansburiana* is characterized by male throat color polymorphism associated with different reproductive strategies (13). Subspecies have been described based on variation in scale traits (14), but there is no reported geographical differentiation in ecomorphological traits (e.g. body size or limbs (15)) despite spanning vastly different environments. Cryptic genetic diversity has been suggested based on mitochondrial data (16–18). Home range, display behaviors, and sexual dimorphism vary throughout its distribution (16, 19, 20), revealing intraspecific variability that does not manifest in species-level differentiation.

We assessed the patterns of differentiation in *U. stansburiana* populations from the geologically/climatically heterogeneous Baja California peninsula, which spans a highly reported deep mitochondrial genetic break (Figure 1A; (17, 18)). The peninsula is a dynamic landscape in which multiple organisms share a co-divergence signal (21, 22).

**Figure 1.**
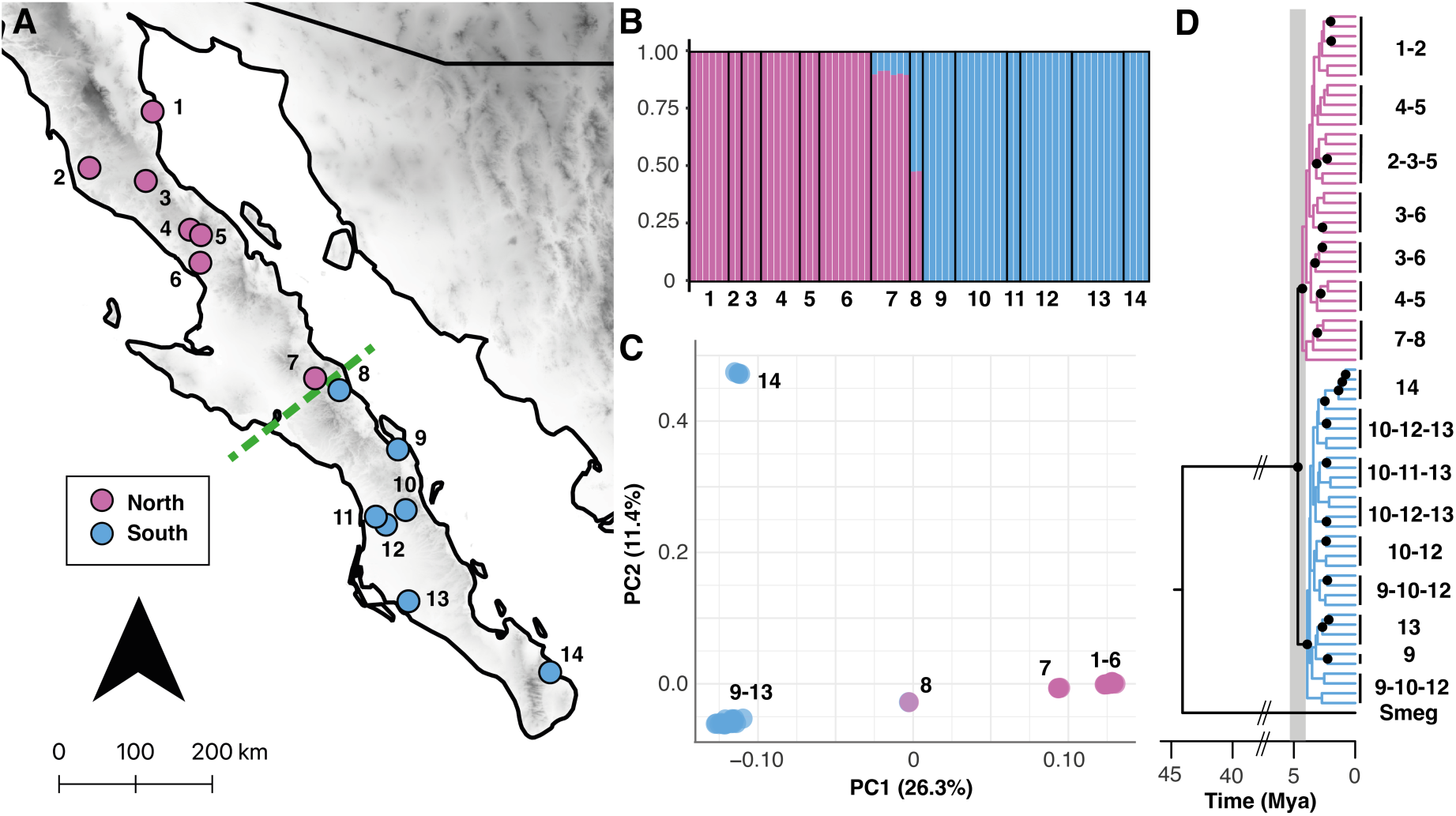
A. Sampling locations for *U. stansburiana* across the Baja California peninsula showing the previously described genetic break by (18). Details for sampling locations are in Table S1. **B.** Bar plot showing the admixture proportions for all individuals. **C.** PCA showing the patterns of genomic variation. **D.** Time-calibrated Bayesian phylogenetic inference. Black-filled circles indicate nodes with a posterior probability higher than 0.9. The gray shaded area represents the 90% posterior density for the node age of (4.01–5.17 Mya).

Previous studies proposed that a mid-peninsular seaway caused population differentiation (18), but this hypothesis has recently been conclusively refuted by detailed geological field mapping, analysis and geochronology (23). Ecological niche models support that populations of *U. stansburiana* were in contact during the Last Glacial Maximum (LGM) and interglacial periods, suggesting that isolation in glacial refugia is also unlikely to explain population divergence (24, 25). Divergence due to stochastic isolation by distance remains a viable hypothesis (26), in which case we would expect to find limited evidence of differential adaptation in association with ecological disparities. In contrast, differential ecological or temporal adaptation driven by monsoon-based rainfall asynchrony could cause or strengthen the observed divergence (21, 22). Here, we used WGS, morphological, habitat use, and behavioral data in integrative statistical frameworks to test which factors are promoting or constraining differentiation among populations of *U. stansburiana* on the Baja California peninsula.

## Results

### Two genetic groups on the Baja California peninsula

Our WGS yielded 5.8x mean coverage per individual (range: 3.8–12.8x). After recalibration and filtering, 10,973,032 genetic variants were retained. Admixture analysis, PCA, phylogenetic reconstruction, and an effective migration surface showed two genetic groups located north and south of the mid-peninsula as the most likely number of clusters, consistent with prior studies (Figure 1B-C; Figure S1; Figure S2; Table S1). The average genome-wide F_ST_ between groups was 0.26 (median = 0.24), and our time-calibrated phylogenetic reconstruction estimated divergence at 4.67 Mya (95% HPD of 4.01–5.17 Mya; Figure 1D).

### Morphological and habitat use are constrained between genetic groups

No significant differences were detected between groups for any ecomorphological trait (Table S2, Figure 2A). However, a significant effect of sex was detected (Table S2, Figure 2A). Males had longer snout-vent lengths (SVL), larger head dimensions (HL and HW), longer limbs (AL, FL) and tails (TL), whereas females had longer trunks (AGD, Figure 2A). Micro-habitat availability differed significantly among groups (Figure S3; Table S3); northern sites were characterized by higher shrub cover, whereas southern locations had higher tree cover. Nonetheless, lizards from all populations showed a significant preference for ground or rock microhabitats when taking habitat availability into account (Table S3, Figure 2B).

**Figure 2.**
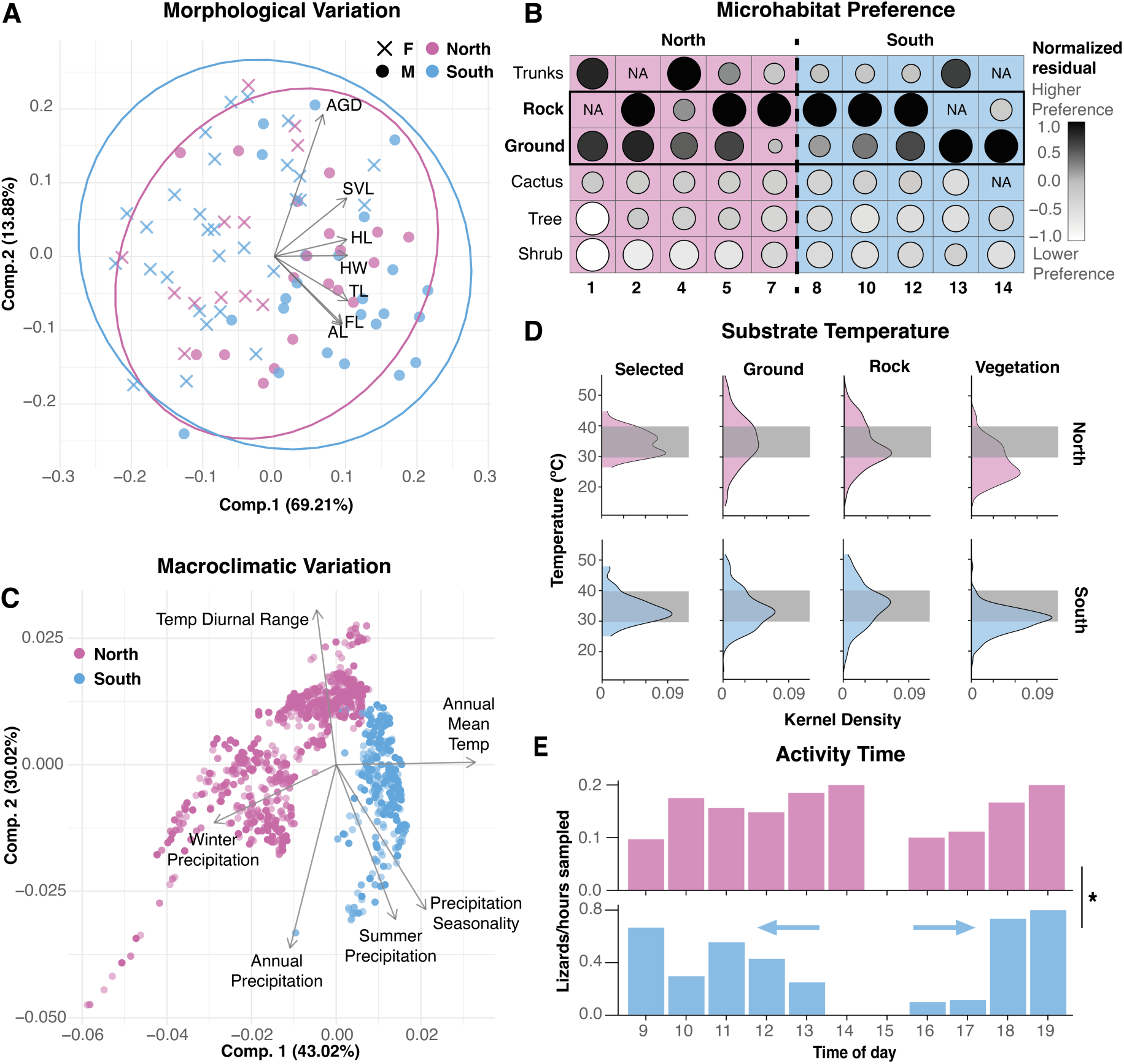
A. PCA showing the lack of differentiation in the morphological variables from both *U. stansburiana* genetic groups. **B.** Microhabitat preference of side-blotched lizards estimated by comparing habitat use versus habitat availability for each location separately. Color and circle size represent the normalized Fisher’s exact test residual. **C.** PCA of 19 bioclimatic variables for *U. stansburiana* on the Baja California peninsula. Table S4 shows the relative contribution of each variable to the PCs. The climatic niche was significantly divergent between northern and southern genetic groups. **D.** Kernel density of temperature measurements for the lizard’s selected substrate and available temperature in different substrate types for northern and southern locations. The gray area represents the 90% density of lizard’s selected temperatures. **E.** Activity time for each genetic group, measured as the number of captured lizards normalized by our sampling effort per time of day. Asterisks indicate significant differences (α = 0.05).

### Thermoregulatory behavior adjusts to different habitats

The two genomic groups live in significantly different macroclimates (niche overlap = 0.043, *p* < 0.001). The southern clade inhabits warmer environments with higher summer precipitation and higher precipitation seasonality, whereas the northern clade inhabits colder environments with higher winter precipitation and lower precipitation seasonality (Figure 2C, Figure S4, Table S4). They also inhabit significantly different microclimates: air temperature and relative humidity at time of capture were significantly higher at southern populations (Figure S5, Table S5). However, the lizards’ selected perch temperature did not significantly differ between groups, indicating the same thermal preference (Figure 2D, Table S5). Substrate temperatures of ground and rock microhabitats peaked within the 90% preferred temperature interval for the species, in contrast with vegetation (Figure 2D), suggesting lizards’ preference for rock and ground substrates is sufficient for thermoregulation regardless of differences in habitat availability. Importantly, activity times differed significantly between genetic groups (Figure 2E, Table S6), with southern populations showing higher activity in cooler hours.

### Most genetic differentiation is explained by physical distance and precipitation

We assessed the relative contribution of physical distance and topographical resistance (i.e., limits to dispersal by elevation changes), temperature, and precipitation on F_ST_ with three independent approaches. First, we detected significant patterns of isolation by distance (Mantel’s r = 0.79; *p* = 0.001), isolation by topography (Mantel’s r = 0.80; *p* = 0.001), and isolation by precipitation (Mantel’s r = 0.84; *p* = 0.001) and no effect of isolation by temperature (Mantel’s r = -0.03, *p* = 0.50; Figure S6). Second, a Generalized Dissimilarity Model (GDM) showed 87% of the variation in population-level genetic differentiation was explained by physical predictors (distance and resistance) and precipitation (Figure 3A).

**Figure 3.**
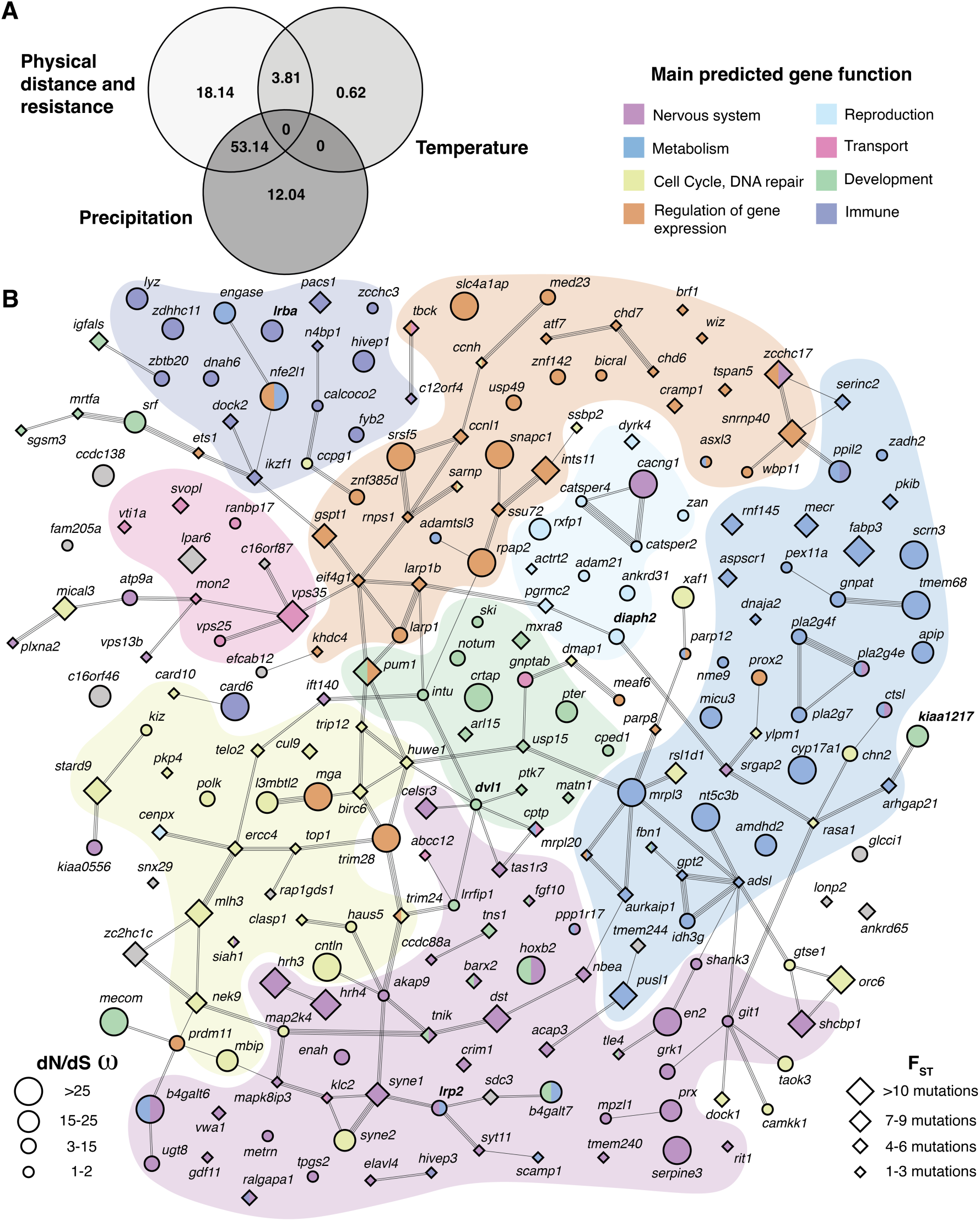
A. Deviance explained by each predictor in the GDM. **B.** String interaction network analysis for genes with potentially functional mutations from the high F_ST_ regions, and genes with μ > 1 and one allele fixed in one of the genetic groups from the dN/dS analysis. Lines represent known interactions between genes, and colors indicate the potential function of each gene. Circle size is proportional to the number of SNPs located in each coding sequence or promoter region for genes from the F_ST_ list, or the μ value for the dN/dS list. Genes overlapping between methodologies are highlighted in bold.

Temperature differences alone contributed < 1% to genetic differentiation (∼ 4% in combination with distance, Figure 3A). Third, a more detailed analysis of Partial Information Decomposition (PID) yielded consistent results: the nodes contributing the highest information about population differentiation (F_ST_) contained distance, resistance, and precipitation (Figure S7). Additionally, distance and resistance were almost entirely redundant (Π = 0.066 bit). About 33% of the information among distance, resistance, and precipitation was redundant, while precipitation was the only variable to provide substantive, unique partial information (Π = 0.018 bit). Temperature contributed almost no information uniquely, redundantly, or synergistically.

### Genomic variation, demography and signatures of ecological adaptation

Nucleotide diversity (π) was higher in the southern group (Figure S8; Figure S9). Both groups showed positive mean Tajima’s D values, but values were higher in the south (Figure S8; Figure S9). Demographic reconstructions show a general trend of a historical decline in effective population size starting at ∼1.25 Mya for both groups and when populations were analyzed separately (Figure S10), and a recent decline and *N_e_* expansion at 10 kya (Figure S11). Southern populations, however, show higher variation in effective population sizes.

We evaluated potential signatures of selection with two approaches. First, we analyzed 63,638 variants located within the highest 1% of F_ST_ windows (F_ST_ cutoff: 0.21), of which 62,627 were SNPs and 911 were indels. Only 476 variants from high F_ST_ regions (0.75%) co-located to potentially functional regions (non-synonymous coding, promoter, and UTR; Figure S12; Table S7). Those putatively functional variants mapped to 131 unique genes (Figure 3B, Table S8). Second, candidate genes were identified by analyzing the proportion of synonymous and non-synonymous polymorphisms (dN/dS) across populations. Here, 113 genes had μ > 1 and had fixed SNPs in at least one of the genetic groups (Figure 3B, Table S9). Most of the genes with μ > 1 (92%) had fixed SNPs in the southern clade (Table S9). Five genes overlapped between both approaches (*dvl1*, *diaph2*, *kiaa1217*, *lrba*, *lrp2*; Figure 3B). Genes from the combined candidate lists had significantly higher interactions than expected by chance (*p* = 0.007). In the protein interaction network, we identified through manual curation eight main functional themes (Figure 3B; Figure S13, Table 1, Table S10).

**Table 1.**
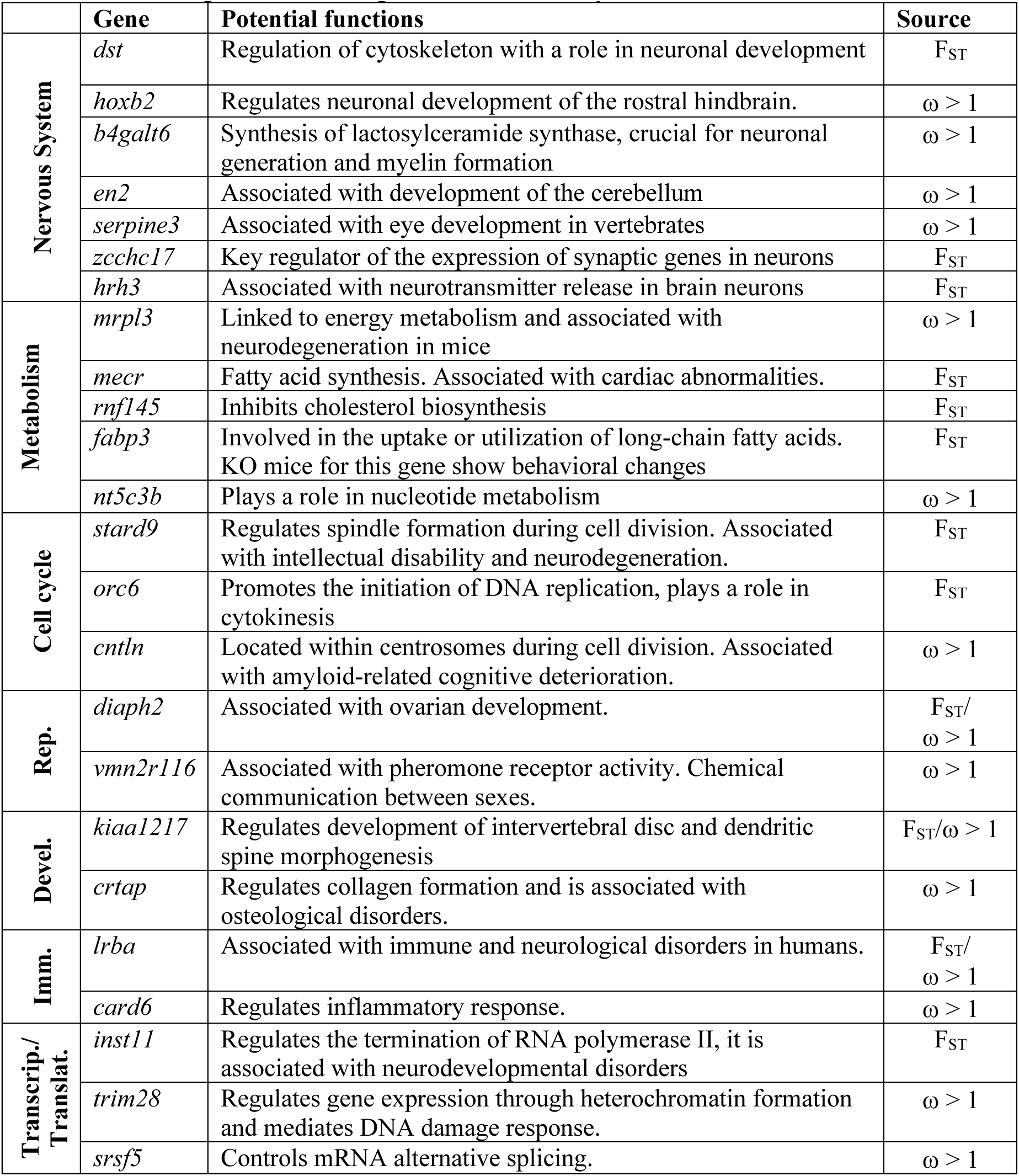
Representative genes and their potential functions, obtained from the candidate lists detected in highest 1% F_ST_ regions or dN/dS analysis. References in Table S11.

Overall, 52 candidate genes were associated with the nervous system (22%), particularly its development (*dst, hoxb2, b4galt6, en2*), neurotransmitter release (*hrh3, zcchc17*), and sensory perception (*grk1*, *serpine3, tas1r3*). We also identified multiple genes associated with biomolecule metabolism (17%; *fabp3, mecr, rnf145*, *nt5c3*), genes related to the regulation of transcription or translation (19%; *ints11, trim28, srsf5*), and cell cycle and DNA repair (16%; *cntln, stard6, orc6*). We also detected some genes relevant for reproduction, associated with pheromone perception (*vmn2r116*), spermatogenesis and ovary development (*actrt2, diaph2*), as well as genes relevant for osteological development (*crtap, kiaa1217*), and immune function (*card6, lrba*). Enrichment analysis identified biological processes associated with the nervous system (e.g. “nervous system development”, “neuron projection morphogenesis”, “neurogenesis”) and biomolecule metabolism (e.g. “metabolic process”, “positive regulation of biosynthetic process”, “positive regulation of cellular metabolic process”) significantly overrepresented in both datasets, despite almost entirely different gene compositions (Table S12).

## Discussion

The study of cryptic species or species with low phenotypic differentiation offers an opportunity to understand mechanisms that can constrain diversification (3, 4). Usually, cryptic lineages occur because of a recent divergence time, convergence, or low divergent selective pressures. Here, we detected high cryptic genetic differentiation in populations of the side-blotched lizard along the Baja California peninsula (Figure 1) and significant climatic and vegetational differences (Figures 2C, S4, S15). Results, however, show a lack of ecomorphological differentiation, which can be explained by decreased *effective* selection pressures due to behavioral adjustments in microhabitat use and activity timing (Figure 2). However, lizards show genomic evidence of adaptation to different rainfall regimes that may be mediated by other traits, such as differences in the nervous system or metabolism (Figure 3, Table 1).

### Drivers of genomic differentiation between U. stansburiana populations

We identified two highly differentiated genetic groups on the Baja California peninsula, matching the previously described mitochondrial genetic structure (17, 18). F_ST_ differentiation was relatively high (0.26) and the estimated divergence age was 4.67 Mya, comparable to species-level divergence reported for other squamates (26, 27). This excludes recent divergence as an explanation for the low phenotypic differentiation.

Genomic variance was largely and redundantly explained by distance and topographic resistance among all populations (Figure 3A, Figure S6, Figure S7). Isolation by distance and resistance are known to generate genetic divergence across the landscape, especially for low-dispersal species (26). Nevertheless, a nearly equal portion of genetic differentiation was explained by precipitation differences (Figure 3A, Figure S6, Figure S7), with much of that information being unique (Figure S7). The Baja California peninsula exhibits a remarkable gradient of precipitation from winter-dominated rainfall in the north to tropically-driven summer monsoons in the south (Figure 2C; Figure S4). Such differences could influence genomic differentiation by allochrony and/or local adaptation (22).

If the species’ phenology responds to seasonal precipitation patterns, this could generate allochrony that reinforces speciation (28). Particularly in arid environments, seasonal precipitation patterns affect ecosystem processes such as the green-up timing of vegetation, which differs here (Figure S14) and resource availability (e.g. abundance of insect prey; (29)), and could drive allochronic speciation as reported for giraffes (30), neotropical frogs (31), birds (32), and desert tortoises (33). Several aspects of the biology of *U. stansburiana* are known to respond to environmental variation in precipitation such as body condition (20), reproductive success (34, 35), and population density and home range size (19). Additionally, the environmental disparity in which *U. stansburiana* populations inhabit along the peninsula (macroclimatic temperature and seasonal precipitation variations, microclimate, and habitat structure) could produce strong divergent selection pressures. Despite using a strict conservative approach to identify candidate genes under selection, we detected genomic signatures of differential adaptation associated with the nervous system, nutrient metabolism, cell cycle, and reproduction.

### Adaptation to different precipitation regimes

Precipitation timing determines the availability of water and food resources in deserts. For instance, arthropod abundance and composition vary seasonally with precipitation on the Baja California peninsula (29). Organisms from arid environments often show signatures of adaptation in osmoregulation processes (36–39), which we did not detect among our candidate genes. As the precipitation differences on the peninsula mostly relate with timing rather than amount, it is not surprising that we detected genes related to signaling and potential response to seasonality of water availability rather than osmoregulatory genes *per se* (Table 1, Figure 3B, Table S12, (33)).

Within our candidate lists, we detected an enrichment of genes associated with the nervous system (Table S12). For instance, *hoxb2*, *dst* and *en2* are associated with its development (40–42); genes that regulate synaptic transmission via neurotransmitter release and receptor density such as *hrh3* and *shank3* differed (43, 44). We also found several genes relevant to sensory perception, such as *serpine3* and *grk1* (associated with eye development (45) and photoreception (46)), as well as perception of sweet taste (*tas1r3*, (47)). Selection on these genes could imply changes in behavior, sensory perception, and mechanisms of response to stress and sporadic sources of water, as proposed for other desert lizards (36), rodents (48) and livestock (49). We also identified an enrichment of genes associated with biomolecule metabolism. For example, the genes *mecr* and *rnf145* associate with fatty acid and cholesterol biosynthesis, respectively, and *fabp3* controls the uptake and utilization of long-chain fatty acids (50). The *nt5c3*, *pusl1* and *nme9* genes are involved in nucleic acid metabolism (51–53). Adaptations in nutrient and energy homeostasis are also commonly reported in genomic studies of desert species, suggesting potential differences in diet, nutrient uptake and storage efficiency following pulses of resource availability (48, 54, 55).

Finally, results showed genes associated with the regulation of DNA transcription (*inst11*, *mga*; (56, 57)), heterochromatin formation (*trim28*; (58)), and mRNA splicing (e.g. *snrnp40*, *srfs5*; (59, 60)). These processes could facilitate the chromatin remodeling and transcriptional response of lizards to seasonal changes, as these are often enacted by regulatory cascades, and have been proposed as a mode of seasonal adaptation in other southwestern reptiles (33). Moreover, we identified genes associated with cell cycle including *stard9* which regulates spindle formation (61), and *orc6* that regulates DNA replication and participates in cytokinesis (62). Additionally, some genes that participate in DNA damage repair such as *trim28* or *mlh3* (58, 63), and apoptosis and cell senescence (*card6* or *rsl1d1* (64, 65)) were detected. Together, these genes could underlie differences in the stress response to UV radiation, as proposed for desert mice, desert tortoises, and deer (48, 66, 67). Importantly, UV radiation is significantly higher in the southern peninsula, suggesting higher selective pressure for these adaptations (Figure S15).

Most candidate genes from the dN/dS analysis were identified as fixed in the southern genetic group (92%, Table S9), suggesting that the conditions in which this clade evolved have promoted efficient adaptation. The southern peninsula receives a strong rainfall contribution from the North American monsoon, which may have strengthened/stabilized after flooding of the Gulf of California ∼6.3 Mya (68–69) and which strengthening during the mid-Pliocene (70). Its strengthening would have exacerbated regional precipitation differences and southern populations potentially adapted to such sporadic water sources. Southern populations experience higher seasonality and more precipitation variability today (Figure S4). Several gene functions in our candidate lists could be related to response to environmental cues (either by the nervous system, metabolism, or regulation of transcription/translation). Similar results have been reported for salmon (71) and thrushes (72) evolving across seasonal gradients. Hence, adaptation to an altered precipitation regime could have amplified the divergence initiated by physical distance/resistance. We interpret these results with some caution, as our divergence window (4.01–5.17 Mya) fits between flooding of the gulf, which offered additional warm water for tropical storms, and leaf wax isotope evidence of mid-Pliocene strengthening of the monsoon ∼3 Ma (70).

### Constraints on phenotypic differentiation between U. stansburiana populations

Despite the high levels of genetic differentiation and strong signatures of selection, the phenotypic divergence was low regarding morphology, habitat use patterns, and temperature adaptation (Figure 2). We propose that consistent thermoregulatory behavior of *U. stansburiana* decreases the effective divergent selection pressure on thermally-relevant morphology and physiology (8). Our results show that despite significant differences in temperature at the macro- and micro-climatic levels, lizards effectively thermoregulated by perch selection and changes in activity time (34, 73, 74), explaining lack of thermal- phenotypic divergence (Figure 3D) (15). It also explains the absence of any signatures of adaptation in genes related to thermoregulation (e.g., vasodilation/vasocontraction, heat shock proteins), as reported for other species inhabiting thermal gradients (75).

## Conclusions

The abiotic causes underlying divergence are often difficult to effectively characterize in natural populations, particularly over landscapes with co-occurring geological and climatic complexity. Here, using the side-blotched lizard from the Baja California peninsula as a model, results show behavioral adjustments to significantly different macro- and micro- climates constrained morphological and genetic thermal adaptation. Instead, isolation by distance and seasonal variation in precipitation explain an exceptional proportion of genetic variation, supported through several analyses, with adaptation in genes related to brain development/function, signaling, and sensory perception. Alternative vicariance and climate refugia hypotheses are not supported. The genetic differentiation in *U. stansburiana* occurs in dozens of diverse taxa, suggesting that the physiography and seasonal precipitation disparity of the peninsula have synergized to produce potentially community-scale divergence on this landscape (22, 26).

## Methodology

### Genomic differentiation among populations

Adult lizards were sampled by noosing across 14 locations across the Baja California peninsula (Collecting permit: 13439/19 DGVS and 03273/20 DGVS, Mexico; CDC import permit 20220221-0651A; and IACUC Protocol: Arizona State University 20-1737R; Figure 3.1). Sampling was done during fall (September/October) and/or spring (March/April) between 8:00 to 20:00 hrs. After capture, morphological and microhabitat measurements were taken for each individual, a tail clip was obtained and preserved in RNAlater, and the lizard was released. Ninety-two lizards were sampled (Table S1). Tissues from 71 samples (Table S1) were sent to Yale Center for Genomic Analysis (YCGA) for DNA extraction, library preparation, and whole genome sequencing. Raw reads were trimmed, filtered for quality and aligned to the *U. stansburiana* reference genome (UCM_Usta_1; (76)). Variant calling and filtering was performed (Supplementary Methods).

To explore the population genetic structure of *U. stansburiana* on the Baja California peninsula we performed a Principal Component Analysis (PCA), and admixture analysis, an effective migration surface, and we estimated the genome-wide window-based Weir and Cockerman’s F_ST_ index between genetic groups (Supplementary Methods).

Furthermore, a Bayesian phylogenetic inference was generated to estimate the divergence age between the two identified genetic groups (Supplementary Methods).

### Phenotypic divergence

#### Morphology

We compared the morphological configuration between genetic groups. For this, we measured the following morphological traits on 85 lizards (including the ones used for genomic analysis; Table S1): body mass, snout-vent length (SVL), axilla-groin distance (AGD), head length (HL), head width (HW), arm length (AL), foot length (FL) and tail length (TL). Patterns of morphological variation were visualized with a PCA and compared between genetic groups in R v4.2.1 (Supplementary Methods, (77)).

#### Microhabitat use patterns

Microhabitat availability was characterized by three 50 m transects randomly placed in each one of 10 sampling locations. The different microhabitat components were categorized as “tree”, “cactus”, “shrub”, “ground”, “rock” or “trunk”, and the relative cover was quantified as the proportion of each component adding the three transects data. Habitat availability was compared among populations. We also recorded lizards’ selected microhabitats in the same locations. For this, one observer registered the perch used by all sighted *U. stansburiana* individuals during the sampling hours. Then, habitat preference was independently evaluated for each location by comparing lizard’s habitat use versus habitat availability.

#### Macroclimatic characterization

Occurrence data for *U. stansburiana* within the Baja California peninsula was obtained. Records were manually curated to represent the species’ distribution and were assigned to each clade based on their geographic location. Data on 19 bioclimatic variables was extracted for each occurrence point. Climatic divergence between clades was assessed by calculating the overlap in the multivariate space of bioclimatic variables between the ellipsoids representing the climatic niche of each clade. Furthermore, climatic niche variation was visualized with a PCA, and the variables that contributed the most to differentiating genetic groups were identified with their loadings in the first two PCs axes (Supplementary Methods).

#### Microclimatic characterization

The preferred microclimatic conditions were also measured for 72 lizards from 11 locations (Table S1). After each lizard capture, the air temperature and relative humidity were registered with a digital thermohygrometer (HTI), and the lizard’s selected substrate temperature was measured in duplicate with an infrared thermometer (Etekcity). Air temperature, relative humidity, and substrate temperature were compared between genetic groups. Additionally, the available temperatures at rock, ground, and vegetation microhabitats were measured by taking 5 random temperature measurements of each microhabitat type within 1 m of the lizard’s capture site. Finally, for each captured lizard we also registered the time of capture. As a proxy to the lizard activity times, we calculated the number of lizards captured in 1 h intervals between 8:00 to 20:00 hrs. To compare the activity times between genetic groups we standardized dividing the number of captured lizards by our sampling effort, quantified as the total number of sampling hours/human for each genetic group (Supplementary Methods).

#### Factors influencing the genomic divergence patterns

We used three independent approaches to test which factors better explained the species’ divergence patterns. We calculated the mean genome-wide weighted F_ST_ between all population pairs. Then, we quantified the physical distance between pairs of populations, and as a measure of the topographic resistance between pairs of populations we estimated the resistance distance using the transition matrix calculated from the digital elevation model of the peninsula obtained from WorldClim. We also calculated the macroclimatic differences between population pairs. We analyzed temperature (bio1 to bio11) and precipitation (bio12 to bio19) variables separately. First, we identified which factors contributed the most to the divergence patterns with a generalized dissimilarity model (GDM; (78)), using the F_ST_ distance matrix as a response variable and the distance, resistance, precipitation and temperature distance matrices as predictors. We calculated the deviance explained by the different predictors from our model categorized by physical (distance and resistance), temperature and precipitation. Second, we used each one of these physical datasets along the pairwise F_ST_ values to test for isolation by distance (IBD), isolation by resistance (IBR) and isolation by thermal and precipitation environments (IBTE, IBPE) using Mantel’s tests. Third, we performed a partial information decomposition (PID) analysis to evaluate the amount of unique, redundant and synergistic information of the different predictor distance matrices (distance, resistance, temperature, precipitation) on F_ST_ (Supplementary Methods, (79)).

#### Genomic features and past demography

For each genetic group we calculated nucleotide diversity (π) along the genome, and Tajima’s D. We also calculated these statistics for both genetic groups together. Then, we reconstructed the past demography of each sampling location and genetic group (Supplementary Methods).

#### Potential signatures of divergent selection

We used two approaches to explore potential signatures of positive selection between the two genetic groups. First, we extracted the variants located within the highest 1% F_ST_ windows. We annotated whether variants were located in coding regions, introns, 5’UTR, 3’UTR, promoter, or other intergenic regions (Supplementary Methods). Further, we identified if the variants located within coding regions corresponded to synonymous or non-synonymous. Second, we calculated the ratio between non-synonymous to synonymous polymorphisms for all the annotated genes among all our samples. Here, a value of μ > 1 indicates a higher polymorphism of non-synonymous versus synonymous sites in a gene (Supplementary Methods). For both candidate gene lists (high F_ST_ and μ > 1) we assessed which Gene Ontology (GO) biological processes were enriched and constructed a gene interaction network to identify groups of genes that interact and perform shared functions (Supplementary Methods).

## Supplementary Methods

### Genomic differentiation among populations

Adult lizards were sampled by noosing across 14 locations across the Baja California peninsula (Collecting permit: 13439/19 DGVS and 03273/20 DGVS, Mexico; CDC import permit 20220221-0651A; and IACUC Protocol: Arizona State University 20-1737R; Figure 3.1). Sampling was done during fall (September/October) and/or spring (March/April) between 8:00 to 20:00 hrs. After capture, morphological and microhabitat measurements were taken for each individual, a tail clip was obtained and preserved in RNAlater, and the lizard was released. Ninety-two lizards were sampled (Table S1). Tissues from 71 samples (Table S1) were sent to Yale Center for Genomic Analysis (YCGA) for DNA extraction, library preparation, and whole genome sequencing in an Illumina Novaseq platform generating 150 bp-long paired-end reads.

Raw reads were trimmed and filtered for quality using fastp v0.23.4 (Chen, 2023). Then, reads were aligned to the *U. stansburiana* reference genome (UCM_Usta_1; (2)) with BWA v0.7.17 (3). Variant calling was performed with GATK 4.5.0.0 (4). Since we studied a non-model organism without previously known genomic variant information, we followed the GATK pipeline recommendations and ran a first round of variant calling without applying base recalibration. Then, we used the uncalibrated variants to guide the base recalibration in a second round of variant calling (5). Variants obtained from this second round were used for further analysis. Variant filtering was performed with VCFtools v0.1.14 (6), considering a minimum quality score of 30 per site, a percentage of missing data lower than 10%, a minimum coverage depth of 6x and a maximum of 30x, and a minor allele frequency of 0.05. Finally, we filtered reads for linkage disequilibrium with Plink v1.9 (7) using a window size of 50 kb, a step of 20 kb, and a r^2^ threshold of 0.2.

To explore the population genetic structure of *U. stansburiana* on the Baja California peninsula a Principal Component Analysis (PCA) was performed with Plink. Then, we exported the eigenvectors and eigenvalues from the PCA and visualized the variation in R v4.2.1 (8). Genetic structure was further explored with admixture v1.3.0 (9). In admixture, we tested for different numbers of genetic clusters (K) from 1 to 5, and we selected the most likely K based on the lowest cross-validation error. Moreover, to geographically visualize the genetic structure among populations, an effective migration surface was estimated with EEMS (10). Since the PCA and admixture suggested a most probable number of clusters of K = 2, we estimated the genome-wide window-based Weir and Cockerman’s F_ST_ index between both genetic groups with VCFtools, using a window size of 100 kb and a step of 50 kb.

A Bayesian phylogenetic inference was generated to estimate the divergence age between the two identified genetic groups. Whole genome sequencing data from one individual of *Sceloporus megalepidurus* (Table S1; (11)) was included as an outgroup. For this analysis, we used the *U. stansburiana* genome annotation to select intergenic regions of 1000 bp length that did not contain repeat elements, and that contained variants within the *U. stansburiana* populations. We generated a “fasta” formatted file for each locus by combining the genome assembly and the “vcf” variant data with vcf2fasta (12). We selected the 68 loci with the highest number of variants including the main 17 scaffolds (representative of chromosomes; (2); Table S13), proportionally to their length. For each locus, the best nucleotide substitution model was identified with Jmodeltest v2 (13) based on the Akaike Information Criterion (AIC). A multilocus phylogenetic tree was generated using BEAST v1.10.4 (14). Three independent MCMC were run for 1,000,000 generations. Runs were combined with tree annotator, considering a burn-in of 100,000 generations. The final consensus tree was time-calibrated with a 44 Mya estimated divergence time between *S. megalepidurus* and *U. stansburiana* (15).

### Phenotypic divergence

#### Morphology

We compared the morphological configuration between genetic groups. For this, we measured the following morphological traits on 85 lizards (including the ones used for genomic analysis; Table S1): body mass, snout-vent length (SVL), axilla-groin distance (AGD), head length (HL), head width (HW), arm length (AL), foot length (FL) and tail length (TL). Since about half of the individuals had an autotomized tail, we used the “imputePCA” function from the “missMDA” library (16) to estimate the missing TL values based on the covariance among variables. Patterns of morphological variation were visualized with a PCA in R. Significant differences for SVL and mass were evaluated with a linear model in R including sex and genetic group as factors, and residual normality was confirmed with a Shapiro-Wilk test. For the other morphological variables, differences were assessed with a MANOVA including sex and genetic group as factors, and SVL as a covariable. Multivariate normality was assessed with an energy test using the ‘energy’ package (17) in R. Since the data did not follow a multivariate normal distribution, a non-parametric permutation MANOVA was run using the ‘adonis2’ function from the ‘vegan’ package (18) in R.

#### Microhabitat use patterns

Microhabitat availability was characterized by three 50 m transects randomly placed in each one of 10 sampling locations. The different microhabitat components were categorized as “tree”, “cactus”, “shrub”, “ground”, “rock” or “trunk”, and the relative cover was quantified as the proportion of each component adding the three transects data. Habitat availability was compared among populations with a Fisher’s exact test in R. We also recorded lizards’ selected microhabitats in the same locations. For this, one observer registered the perch used by all sighted *U. stansburiana* individuals during the sampling hours. Then, habitat preference was independently evaluated for each location by comparing lizard’s habitat use versus habitat availability with a Fisher’s exact test in R.

#### Macroclimatic characterization

Occurrence data for *U. stansburiana* within the Baja California peninsula was obtained from the Global Biodiversity Information Facility (19). Records were manually curated to represent the species’ distribution. Data on the 19 bioclimatic variables from WorldClim v2.1 (20) was extracted for each occurrence point with the point sampling tool complement in QGIS v3.16 (21). Occurrence records were assigned to each clade based on their geographic location. Climatic divergence between clades was assessed by calculating the overlap in the multivariate space of bioclimatic variables between the ellipsoids representing the climatic niche of each clade, with the permutation-based test from the ‘ellipsenm’ package (22) in R. Furthermore, climatic niche variation was visualized with a PCA, and the variables that contributed the most to differentiating genetic groups were identified with their loadings in the first two PC axes.

#### Microclimatic characterization

The preferred microclimatic conditions were also measured for 72 lizards from 11 locations (Table S1). After each lizard capture, the air temperature and relative humidity were registered with a digital thermohygrometer (HTI), and the lizard’s selected substrate temperature was measured in duplicate with an infrared thermometer (Etekcity). Air temperature, relative humidity, and substrate temperature were compared between genetic groups with a linear model in R, and residual normality was checked with a Shapiro-Wilk test. Additionally, the available temperatures at rock, ground, and vegetation microhabitats were measured by taking 5 random temperature measurements of each microhabitat type within 1 m of the lizard’s capture site. Finally, for each captured lizard we also registered the time of capture. As a proxy to the lizard activity times, we calculated the number of lizards captured in 1 h intervals between 8:00 to 20:00 hrs. To compare the activity times between genetic groups we standardized dividing the number of captured lizards by our sampling effort, quantified as the total number of sampling hours/human for each genetic group. Differences between the activity time frequency distribution were assessed with a Kolmogorov-Smirnov test in R.

#### Factors influencing the genomic divergence patterns

We used two independent approaches to test which factors better explained the species’ divergence patterns. First, we calculated the mean genome-wide weighted F_ST_ between all population pairs with vcftools. Then, we quantified the physical distance between pairs of populations in R with the ‘gdistance’ package (23). As a measure of the topographic resistance between pairs of populations, we estimated the resistance distance using the transition matrix calculated from the digital elevation model of the peninsula obtained from WorldClim, with the ‘commute distance’ function from ‘gdistance’. We also calculated the macroclimatic differences between population pairs. We analyzed temperature (bio1 to bio11) and precipitation (bio12 to bio19) variables separately. For each set, we calculated the variance-covariance matrix of the scaled and centered variables obtained from our filtered occurrence dataset. Then, we calculated the Mahalanobis distances for all population pairs in the respective temperature and precipitation multivariate spaces. Finally, we used each one of these physical datasets along the pairwise F_ST_ values to test for isolation by distance (IBD), isolation by resistance (IBR) and isolation by thermal and precipitation environments (IBTE, IBPE) using Mantel’s tests in R. To identify which factors contributed the most to the divergence patterns, a generalized dissimilarity model (GDM; Mokany et al., 2022) was performed using the F_ST_ distance matrix as a response variable and the distance, resistance, precipitation and temperature distance matrices as predictors in R using the ‘gdm’ library (25). Then, we calculated the deviance explained by the different predictors from our model categorized by physical (distance and resistance), temperature and precipitation.

Second, we performed a partial information decomposition (PID) analysis to evaluate the amount of unique, redundant and synergistic information of the different predictor distance matrices (distance, resistance, temperature, precipitation) on F_ST_. Information decomposition analyses have been shown useful to disentangling the effect of multiple predictors on complex biological systems (26). We employed the I_min_ formalism of PID (27) because while alternatives exist (28), there are no *a priori* criteria with which to select between them, nor is there unambiguous evidence that any one formalism is broadly superior (29). Furthermore, the I_min_ formalism is relatively simple and was sufficient and intuitive for comparing with gdm. The I_min_ formalism requires discrete data, so each of the variables was independently binned using the nonparametric Bayesian block model (30) as implemented in the ‘Discretizers.jl’ package (v3.2.3, (31)) with F_ST_ binned into 5 bins, resistance and temperature into 3 bins each, and distance and precipitation into 2 bins each. We note that binning approach could affect results. A PID lattice describing the I_min_ decomposition of the information that the predictors (distance, resistance, temperature, precipitation) provide about F_ST_ was computed using the ‘Imogen.jl’ package (v0.4.0, (32)), resulting in 166 nodes. Each node represents the contribution of some combination of variables, the partial information (Π) corresponds to the amount of information that each node contributes, and I_min_ is the sum of each node’s partial information and all nodes below it. For easier representation, we discarded nodes with partial information less than 0.01 bit.

#### Genomic features and past demography

For each genetic group we used vcftools to calculate nucleotide diversity (ρε) along the genome using a window size of 100 kb and a step of 50 kb, and Tajima’s D in non- overlapping windows of 100 kb. We also calculated these statistics for both genetic groups together. Then, we reconstructed the past demography of each sampling location and genetic group by implementing the software SMC++ (33), and the site frequency spectrum (SFS) based approach Stairwayplot2 (34). For both approaches, we used a mutation rate of 2.1\**e*^-10^(35) and a generation time of 1 year (36).

#### Potential signatures of divergent selection

We used two approaches to explore potential signatures of positive selection between the two genetic groups. First, we extracted the variants located within the highest 1% F_ST_ windows. We annotated all SNPs and indels from this set with the ‘locateVariants’ function from the ‘VariantAnnotation’ package (37) in R, identifying whether variants were located in coding regions, introns, 5’UTR, 3’UTR, within the 1,000 bp upstream genes (potentially the promoter, Andersson & Sandelin, 2020), or other intergenic regions. Further, we identified if the variants located within coding regions corresponded to synonymous or non- synonymous. Second, we calculated the ratio between non-synonymous to synonymous polymorphisms for all the annotated genes among all our samples using genomegamap v1.0 (39). Here, a value of μ > 1 indicates a higher polymorphism of non-synonymous versus synonymous sites in a gene. We identified genes with μ > 1 as our candidate gene list. Then, we filtered our candidate genes by the allele frequencies of variants across genetic groups, retaining those genes with at least one non-synonymous allele fixed (p or q = 1) in one of the genetic groups. A less restrictive filtering threshold (p or q > 0.85) was also explored to identify potentially adaptive genes being excluded. For both candidate gene lists (high F_ST_ and μ > 1) we assessed which Gene Ontology (GO) biological processes were enriched with an enrichment analysis performed in g:Profiler ve111_eg58_p18_30541362 (40) using the functional annotations from *Anolis carolinensis* and mouse as references, and a Benjamini-Hochberg’s false discovery rate (FDR) correction for multiple comparisons. Finally, we used STRING v12.0 (41) to construct a gene interaction network for the genes from each gene list together (high F_ST_ and μ > 1) to identify groups of genes that interact and perform shared functions. Two researchers independently classified genes according to their known functions based on the GeneCards (42) and Uniprot (43) databases. When conflicted, the functions were revised, and a consensus was drawn.

---

**Figure S1.**
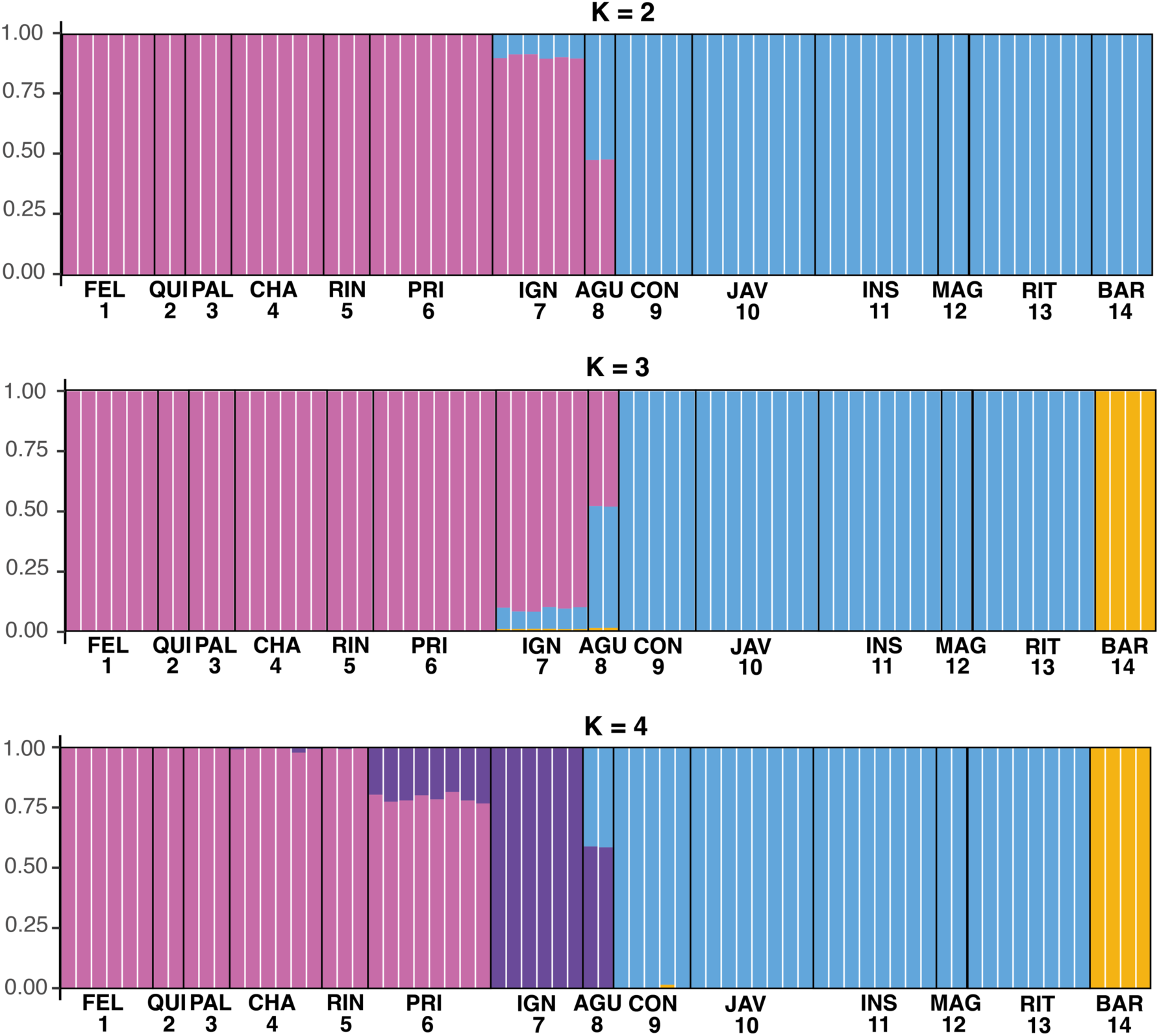
Structure plots for K = 2 to K = 4 for *Uta stansburiana* populations on the Baja California peninsula.

**Figure S2.**
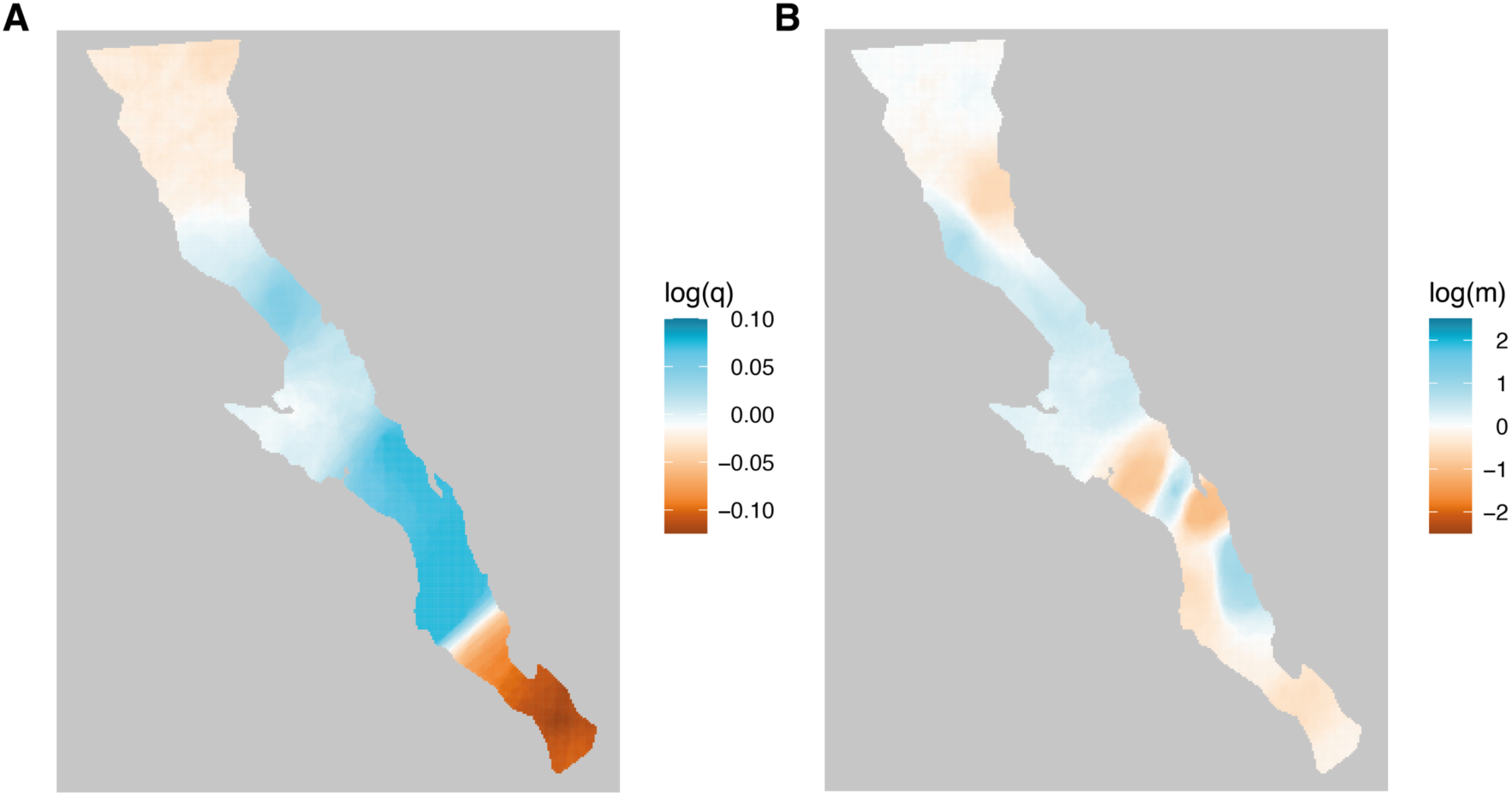
Effective genetic diversity (**A**) and migration (**B**) surfaces for Uta stansburiana on the Baja California peninsula.

**Figure S3.**
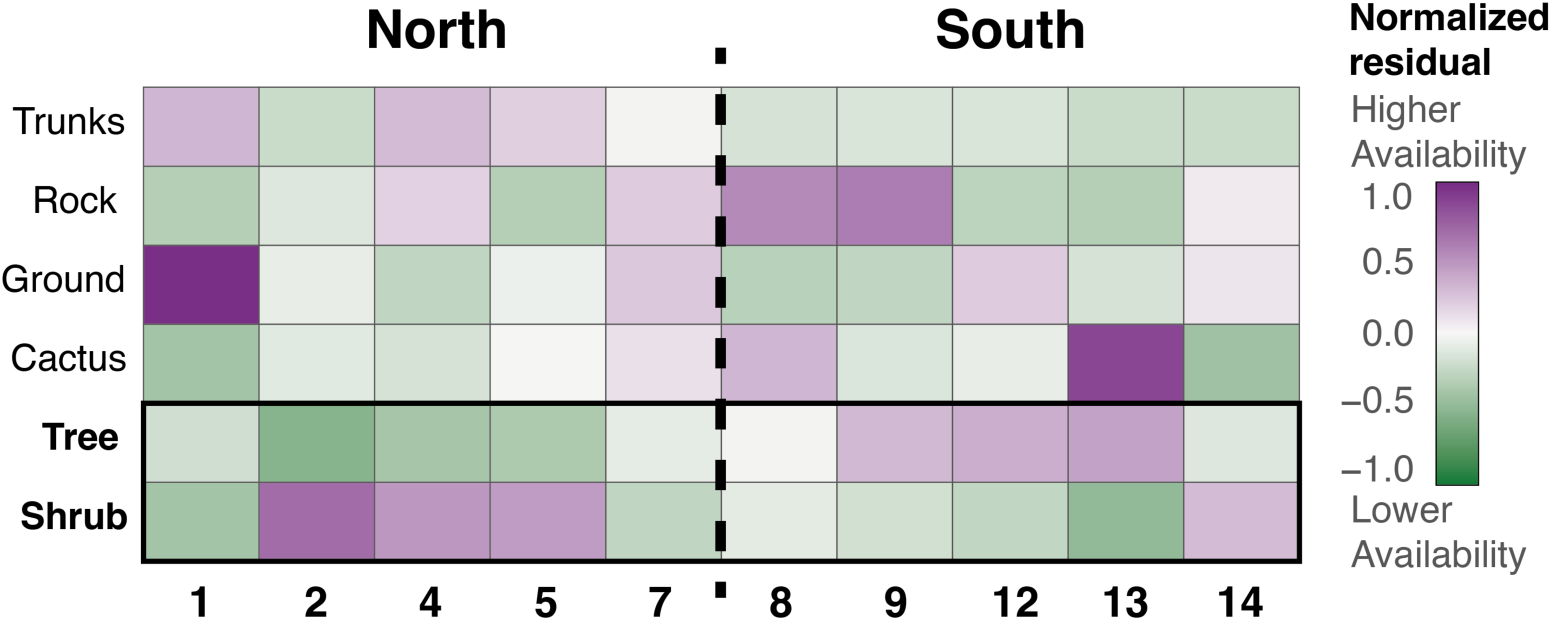
Habitat availability for ten sampling locations of *Uta stansburiana* on the Baja California peninsula. In general, northern sites have higher shrub cover whereas southern sites have higher tree cover.

**Figure S4.**
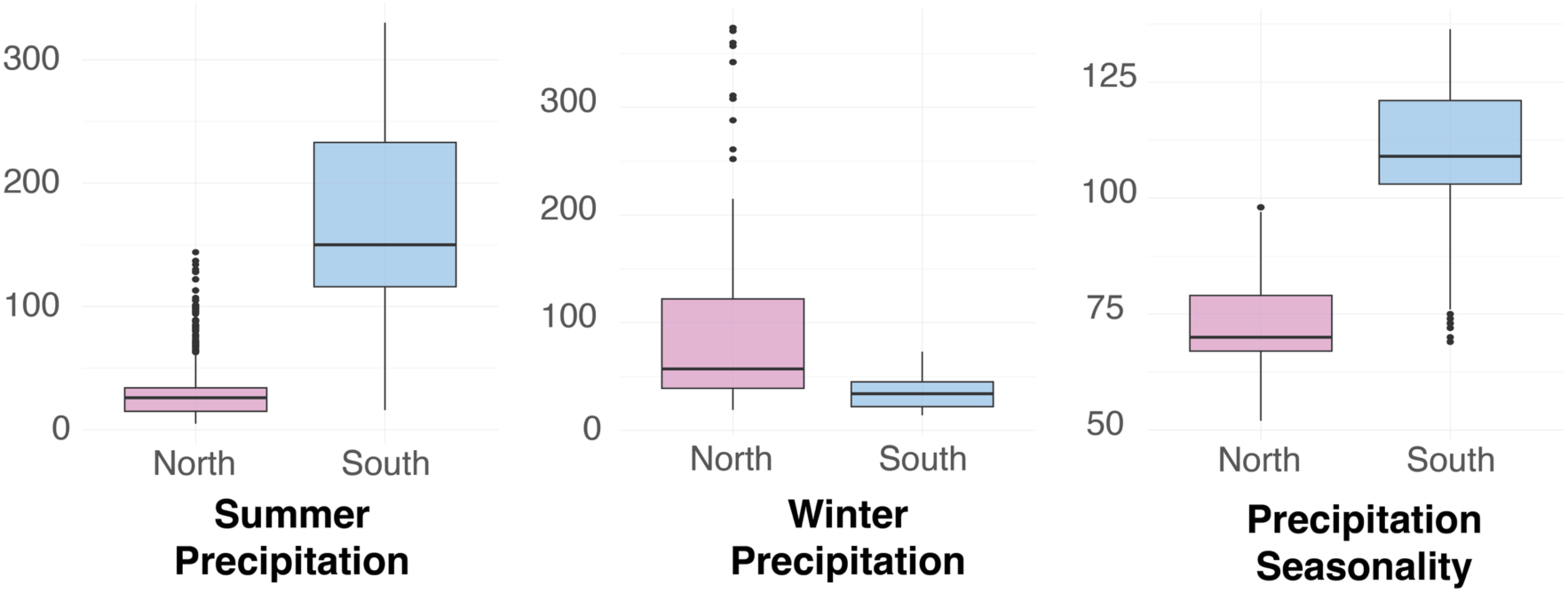
Variation in precipitation variables (mm) for the northern and southern locations on the Baja California peninsula.

**Figure S5.**
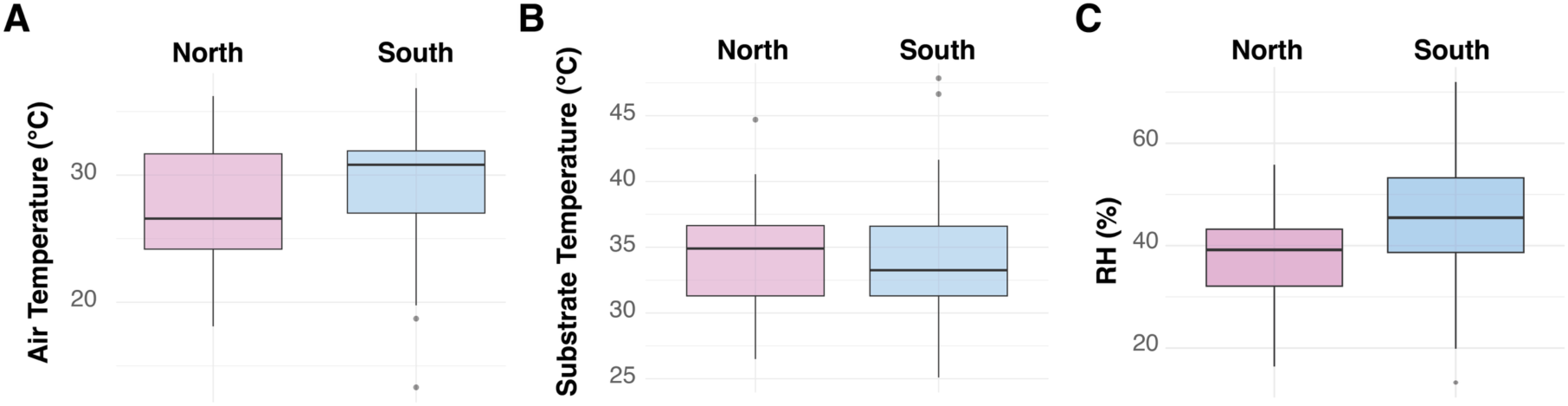
A. Air temperature, **B.** Substrate temperature, and **C.** Ambient relative humidity for *Uta stansburiana* from the northern and southern genetic groups on the Baja California peninsula.

**Figure S6.**
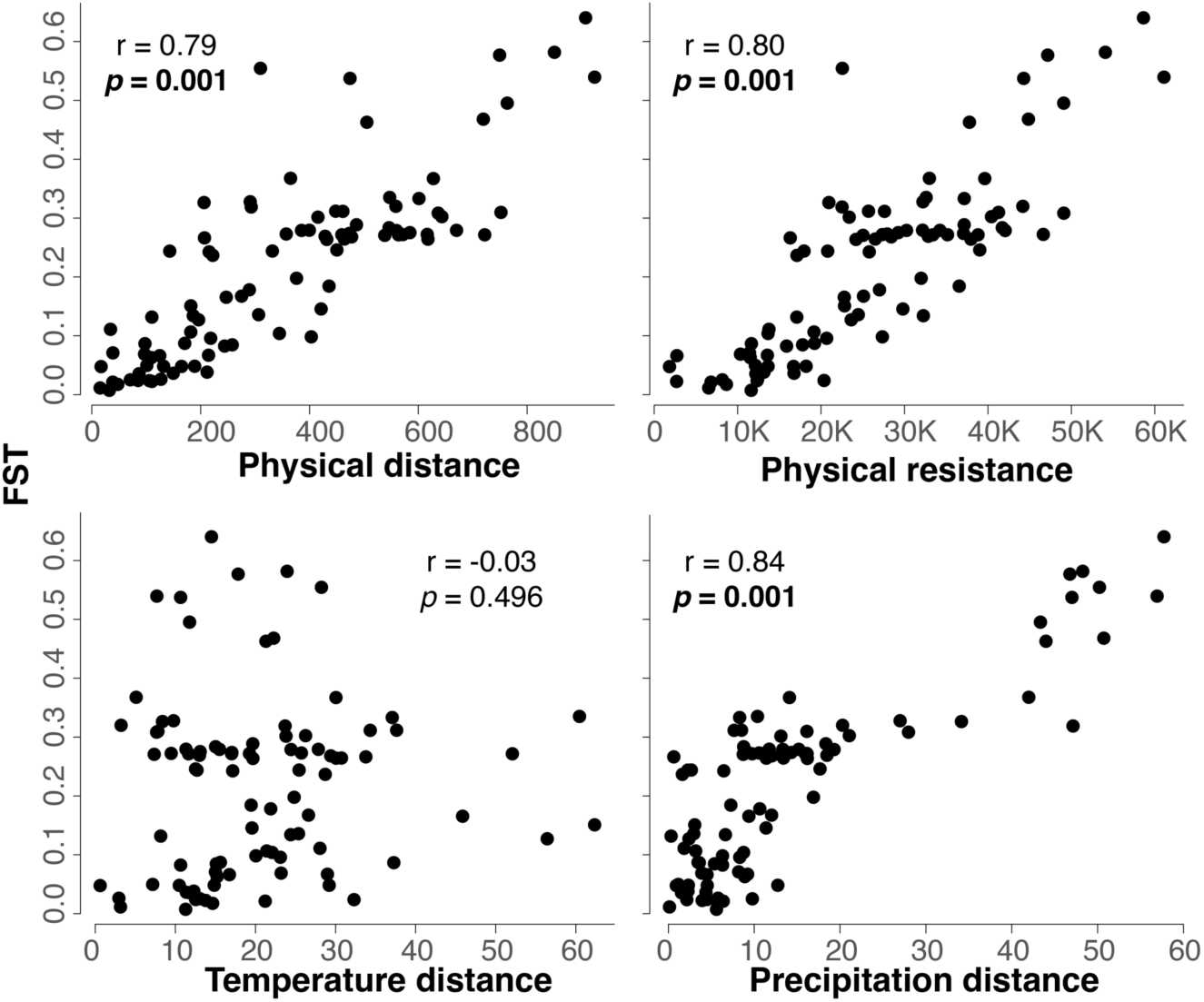
Mantel tests for isolation by distance, isolation by topographic resistance and isolation by temperature and precipitation distance.

**Figure S7.**
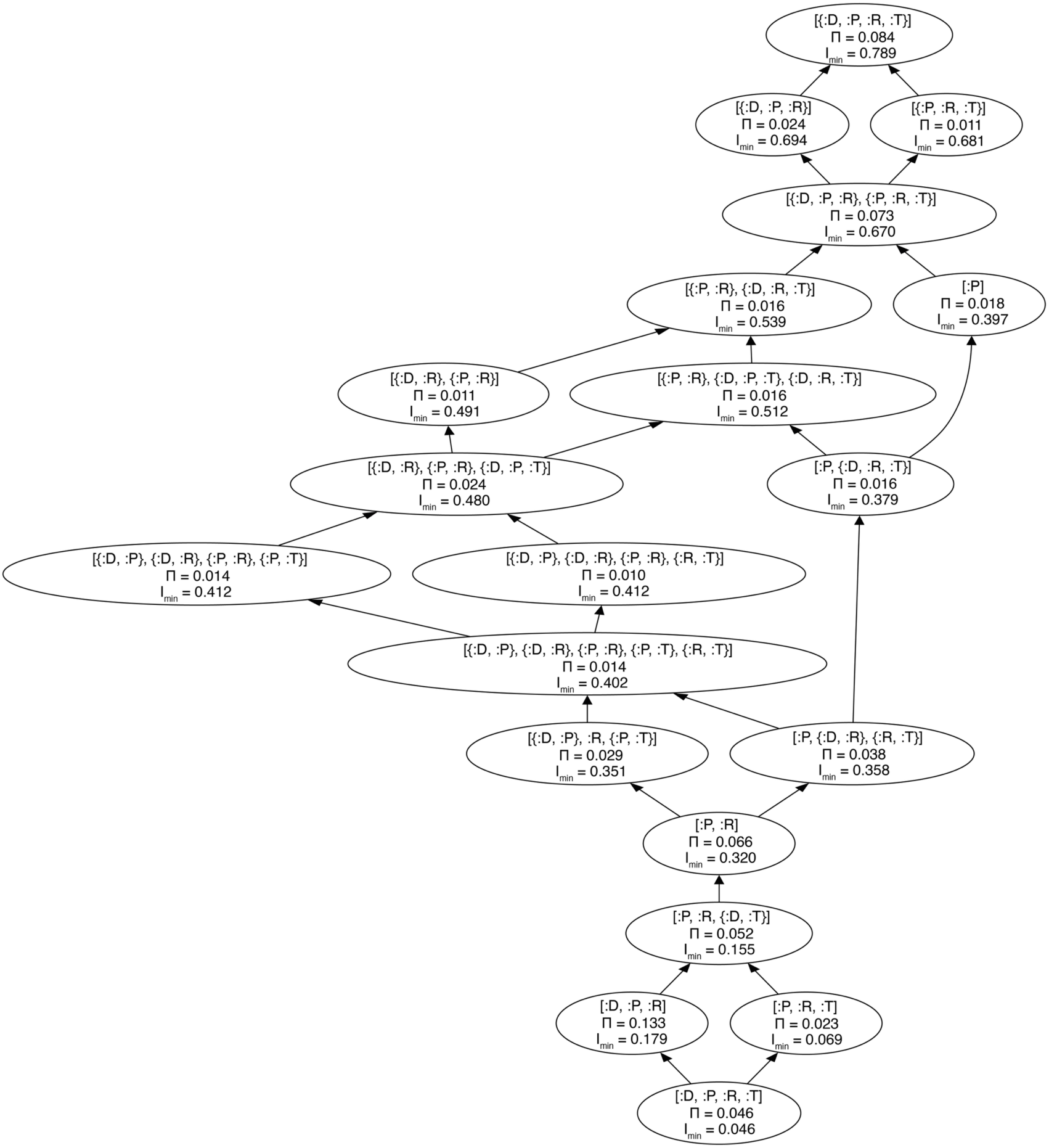
Partial information decomposition lattice for between-population F_ST_ for Uta stansburiana. Each node in the lattice represents the information contribution of a combination of environmental distance predictors (D: distance, R: resistance, T: temperature, P: precipitation). In the lattice, [:X, :Y] represents redundant information between variables X and Y, whereas [{:X, :Y}] represents synergistic information between X and Y.

**Figure S8.**
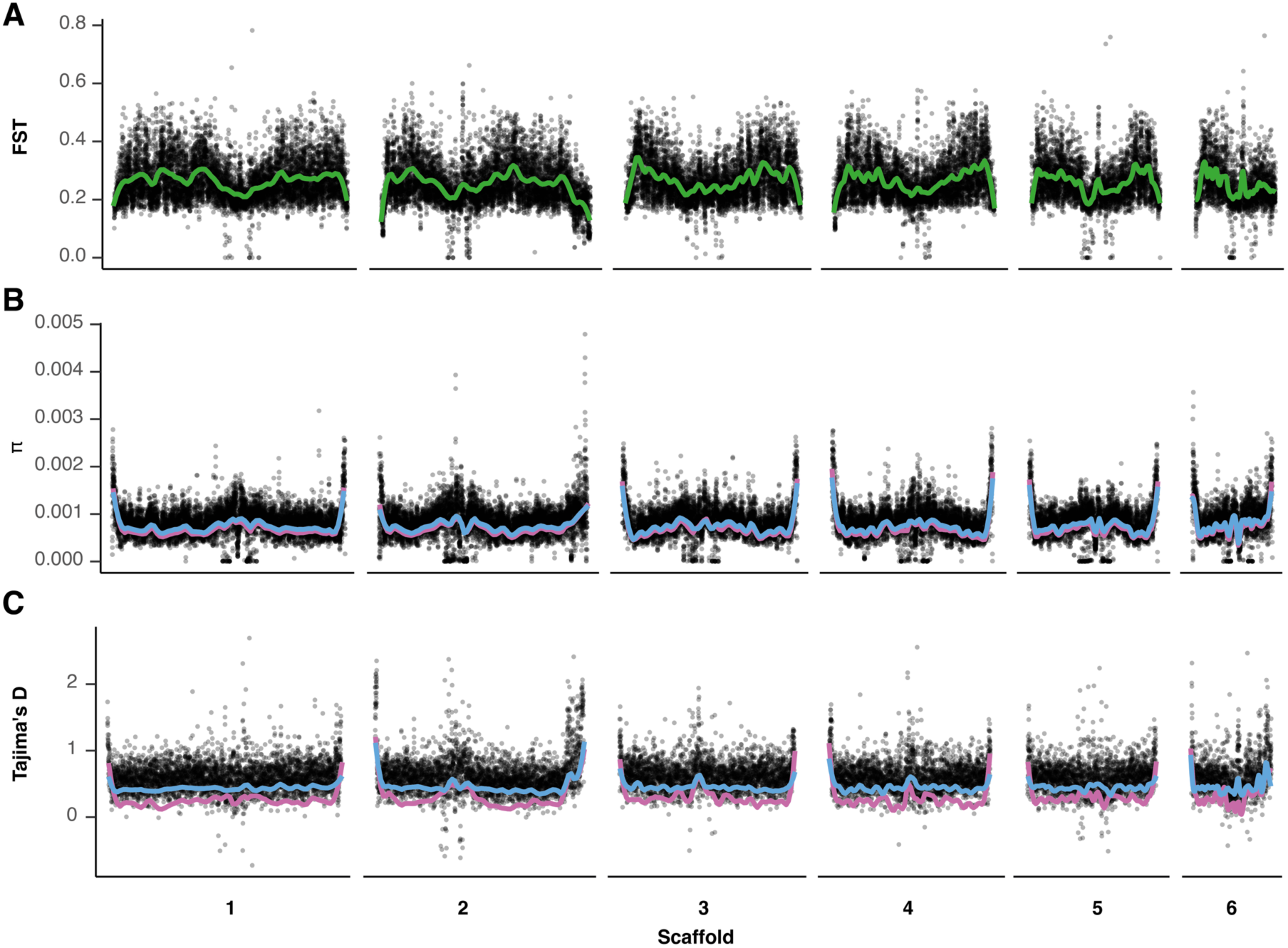
Population genetic statistics calculated along the genome for both genetic groups of *U. stansburiana*. Plot shows values for scaffolds 1-6 corresponding to the macrochromosomes. The northern genetic group is in pink, and the southern in blue. **A.** F_ST_. **B.** π. **C.** Tajima’s D.

**Figure S9.**
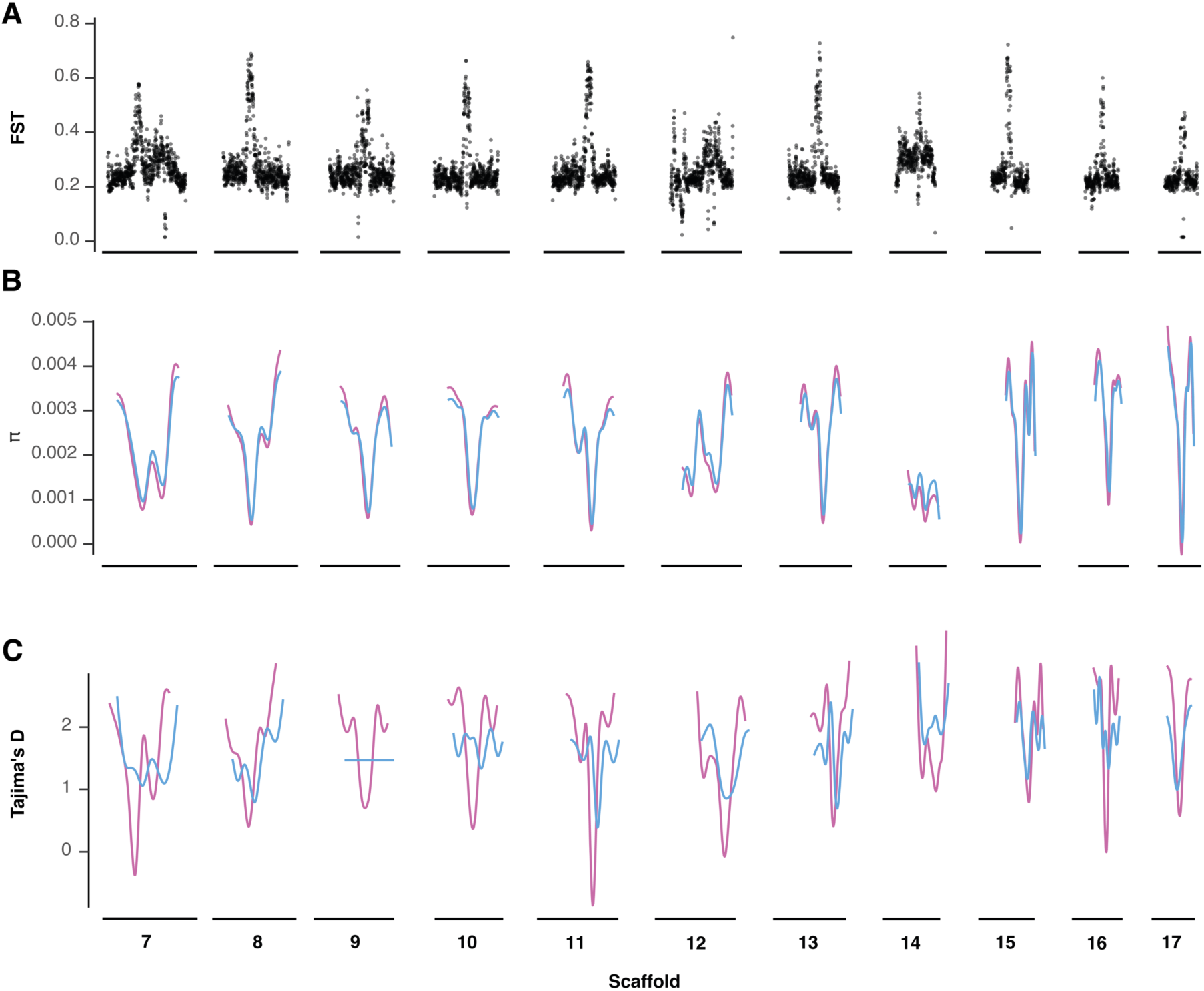
F_ST_, Nucleotide diversity and Tajima’s D for the micro-chromosomes of *Uta stansburiana* populations on the Baja California peninsula. The northern genetic group is in pink, and the southern in blue.

**Figure S10.**
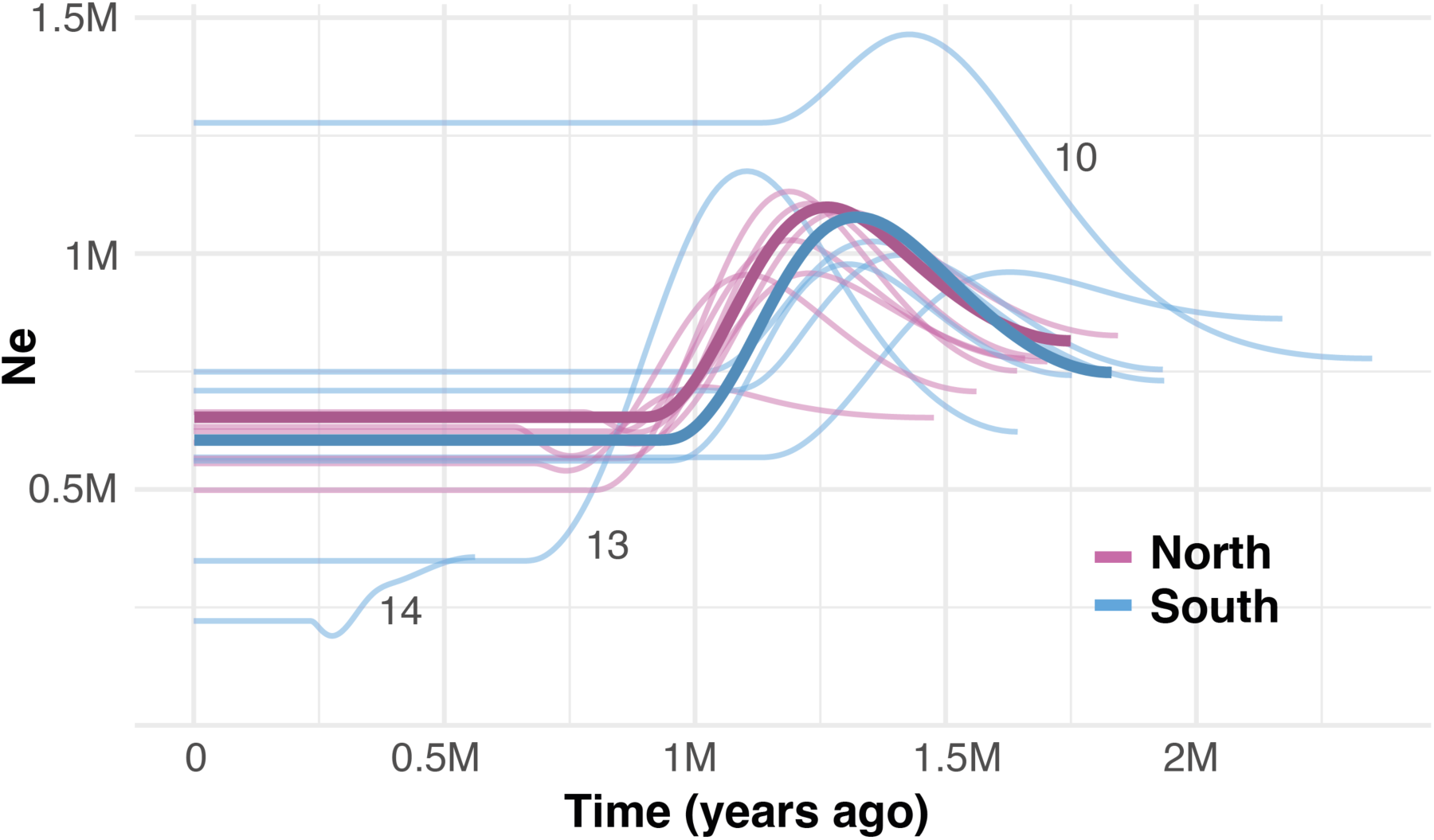
Past demography reconstruction with SMC++ for each population (light colors) and genetic group (darker colors) of U. stansburiana on the Baja California peninsula.

**Figure S11.**
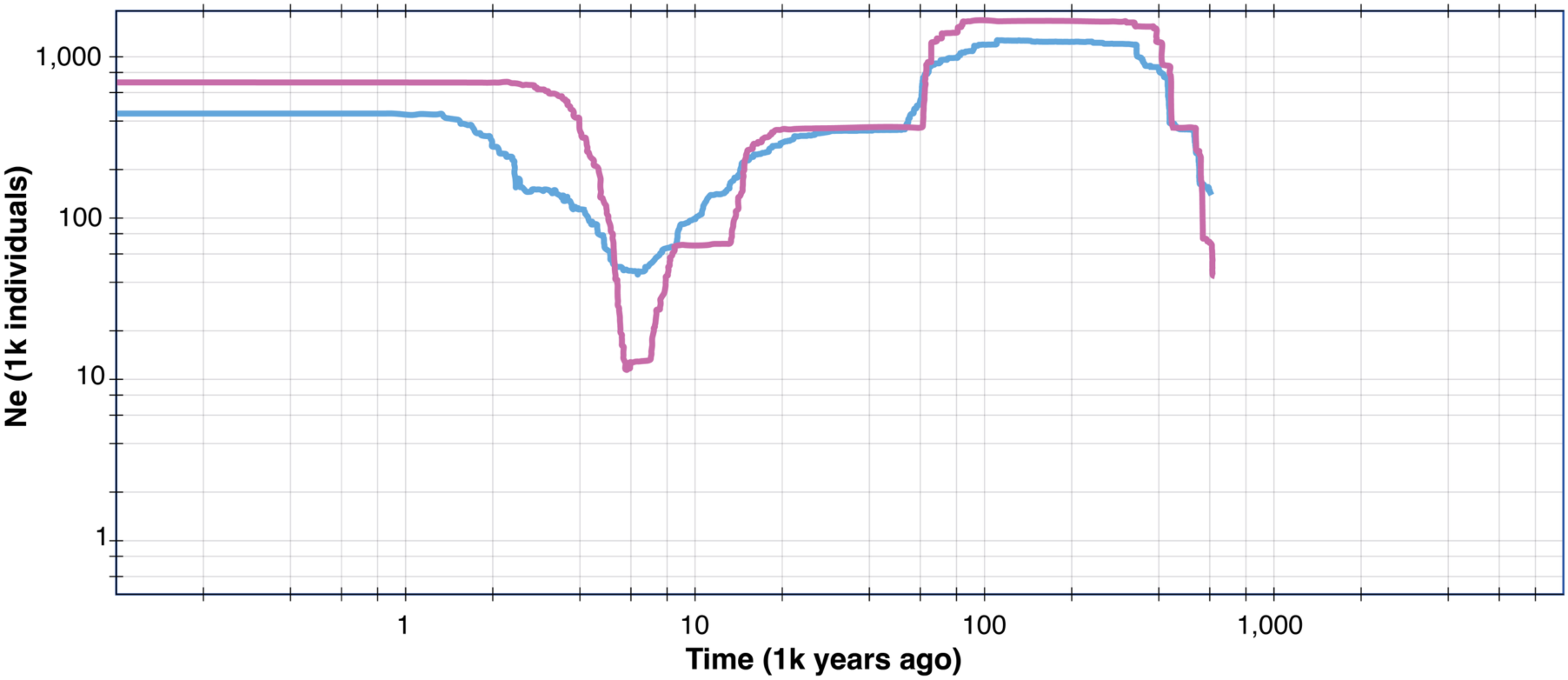
Demographic analysis performed with Stairwayplot on both genetic groups of *Uta stansburiana* on the Baja California peninsula.

**Figure S12.**
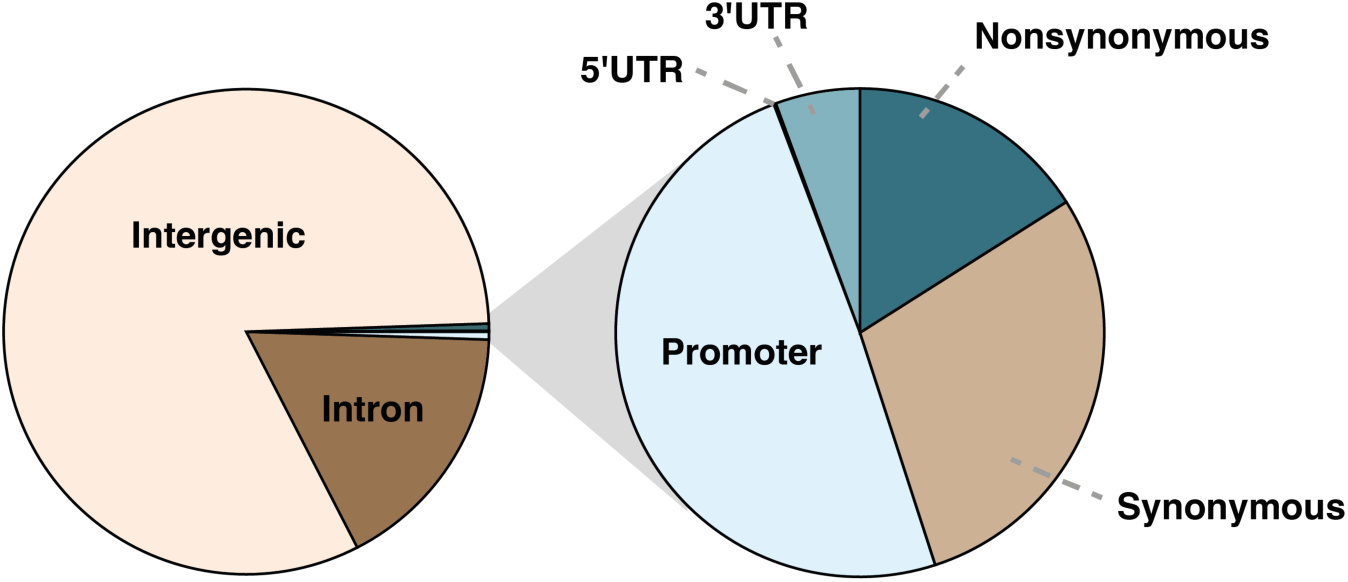
Functional classification of variants located within the highest 1% F_ST_ regions between northern and southern genetic groups of *Uta stansburiana*.

**Figure S13.**
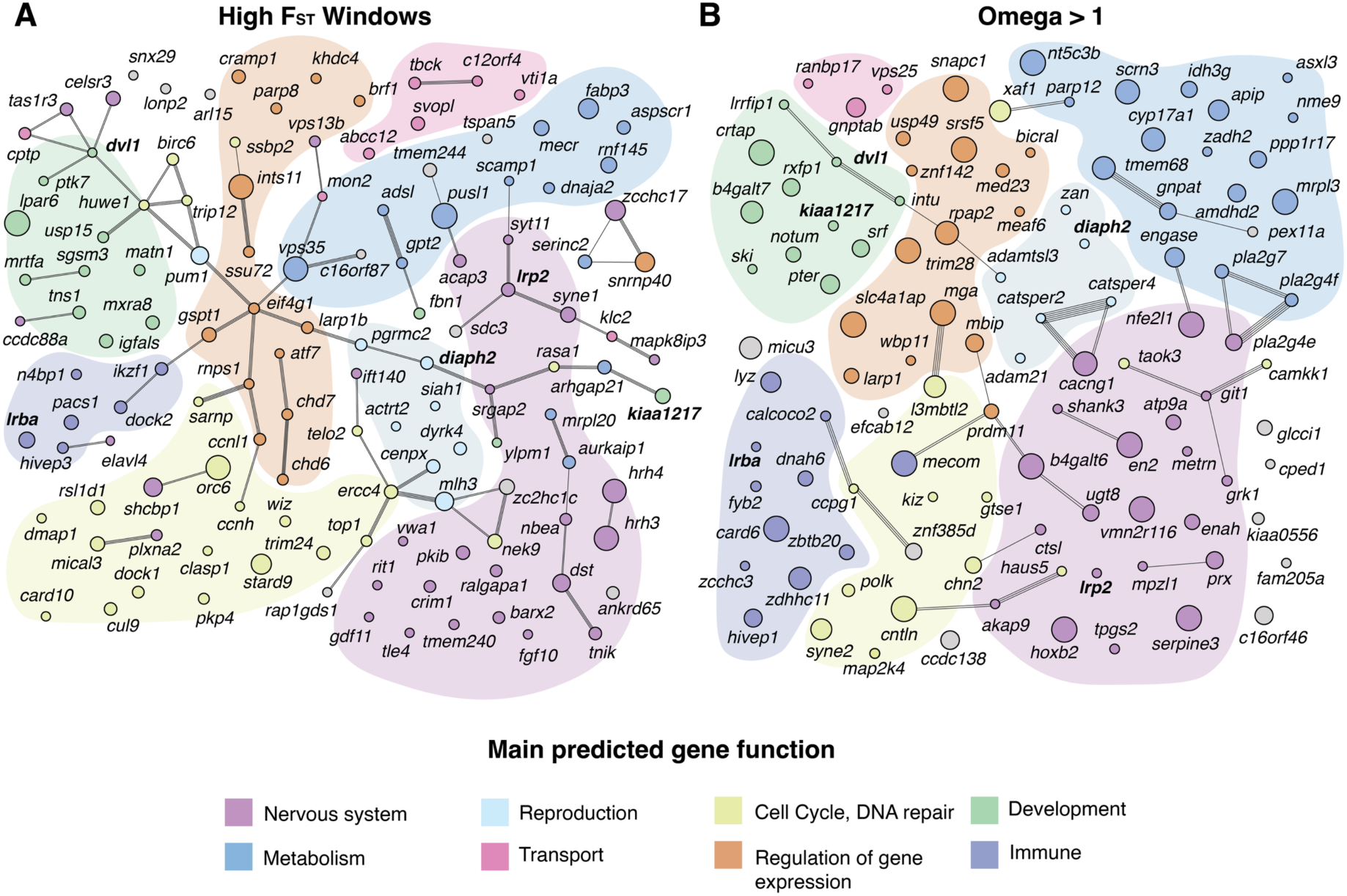
A. String network analysis for genes from the high FST regions with variants located in their promoter region, or non-synonymous variants in their coding regions. **B.** String network analysis for genes with μ > 1, and one allele fixed in one of the genetic groups. Lines represent known interactions between genes, and colors indicate the potential function of each gene. Circle size is proportional to the number of SNPs located in each coding sequence or promoter region in **A**, and the μ value in **B**. Genes overlapping between methodologies are highlighted in bold.

**Figure S14.**
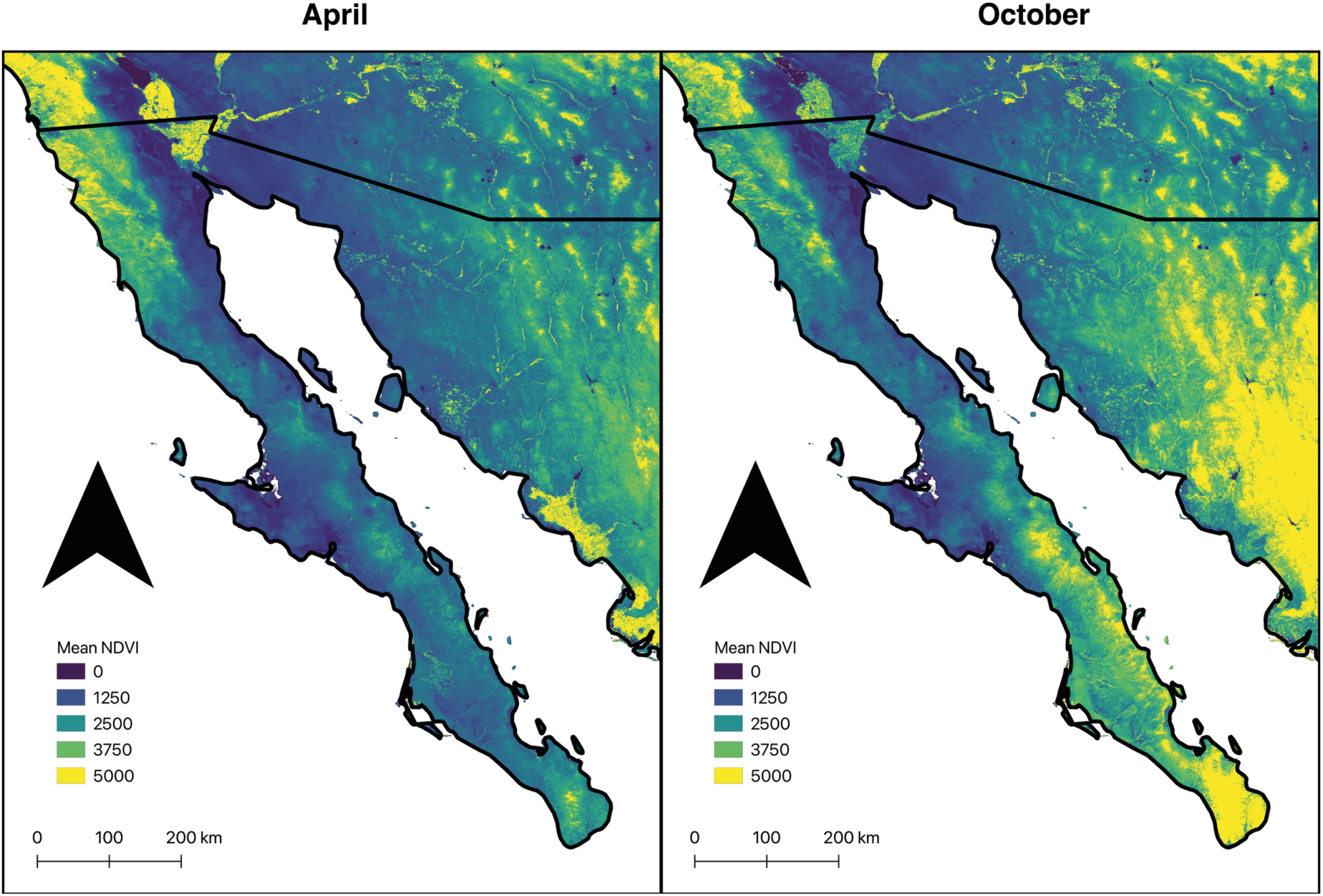
Seasonal variation in vegetation green-up time along the Baja California peninsula showed as the average NDVI values between 1993-2023 for April and October.

**Figure S15.**
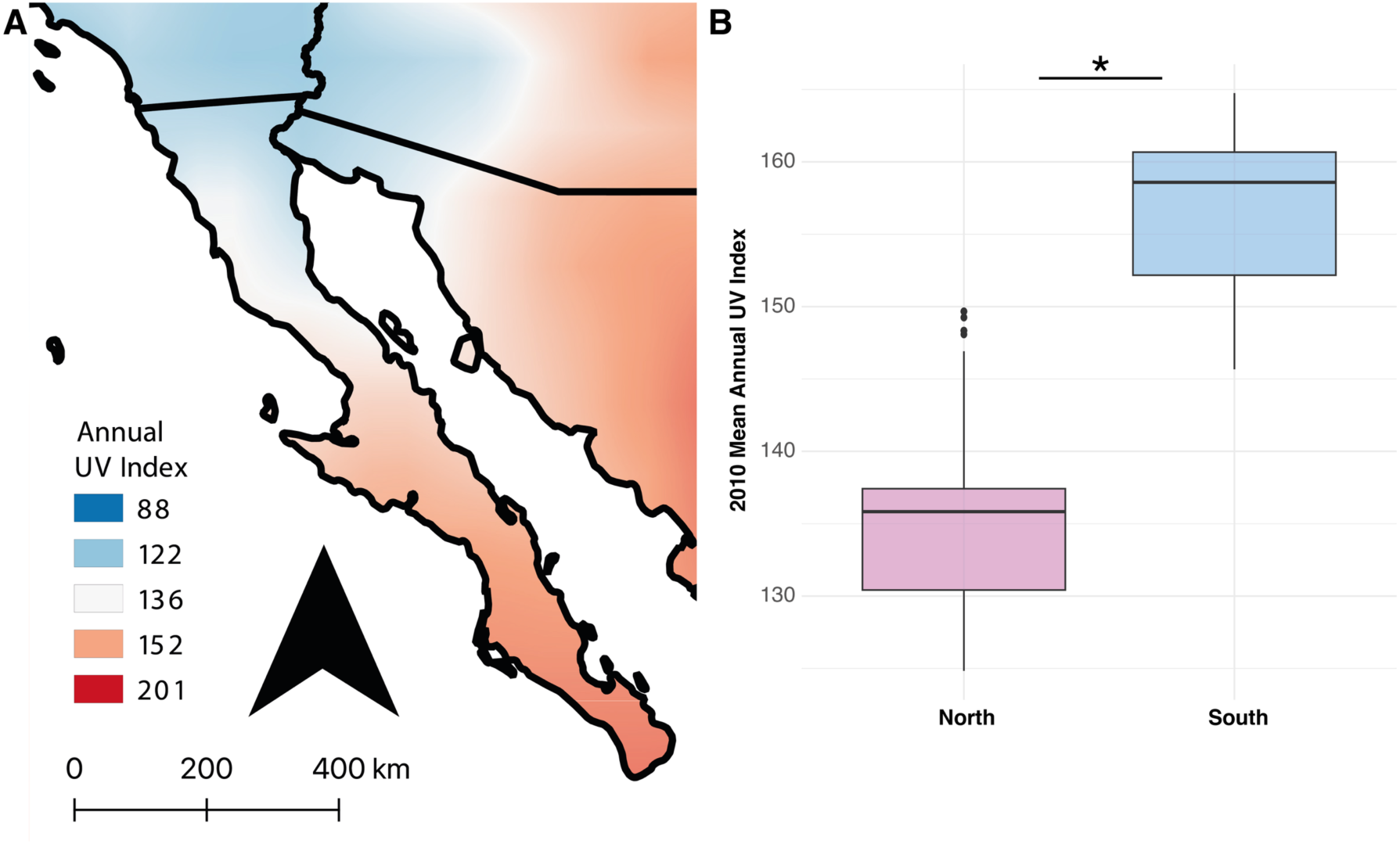
A. Variation of UV radiation across the Baja California peninsula. **B.** The northern and southern regions have significantly (p < 0.001) different annual UV indices.

## References

1. R. De-Kayne, et al., Why Do Some Lineages Radiate While Others Do Not? Perspectives for Future Research on Adaptive Radiations. Cold Spring Harb Perspect Biol a041448 (2024). 10.1101/cshperspect.a041448.

2. C. H. Martin, E. J. Richards, The Paradox Behind the Pattern of Rapid Adaptive Radiation: How Can the Speciation Process Sustain Itself Through an Early Burst? Annu. Rev. Ecol. Evol. Syst. 50, 569–593 (2019).

3. C. Fišer, C. T. Robinson, F. Malard, Cryptic species as a window into the paradigm shift of the species concept. Molecular Ecology 27, 613–635 (2018).

4. T. H. Struck, et al., Finding Evolutionary Processes Hidden in Cryptic Species. Trends in Ecology & Evolution 33, 153–163 (2018).

5. K. Winker, Reuniting Phenotype and Genotype in Biodiversity Research. BioScience 59, 657–665 (2009).

6. D. Bickford, et al., Cryptic species as a window on diversity and conservation. Trends in Ecology & Evolution 22, 148–155 (2007).

7. D. W. Pfennig, et al., Phenotypic plasticity’s impacts on diversification and speciation. Trends in Ecology & Evolution 25, 459–467 (2010).

8. M. M. Muñoz, The Bogert effect, a factor in evolution. Evolution 76, 49–66 (2022).

9. C. R. Campbell, J. W. Poelstra, A. D. Yoder, What is Speciation Genomics? The roles of ecology, gene flow, and genomic architecture in the formation of species. Biological Journal of the Linnean Society 124, 561–583 (2018).

10. D. Schluter, L. H. Rieseberg, Three problems in the genetics of speciation by selection. Proc. Natl. Acad. Sci. U.S.A. 119, e2122153119 (2022).

11. A. D. Leaché, et al., Phylogenomics of Phrynosomatid Lizards: Conflicting Signals from Sequence Capture versus Restriction Site Associated DNA Sequencing. Genome Biology and Evolution 7, 706–719 (2015).

12. P. Uetz, et al., *The Reptile Database*, http://www.reptile-database.org (2024).

13. B. Sinervo, C. M. Lively, The rock–paper–scissors game and the evolution of alternative male strategies. Nature 380, 240–243 (1996).

14. R. E. Ballinger, D. W. Tinkle, Systematics and Evolution of the Genus Uta (Sauria: Iguanidae) (Museum of Zoology, University of Michigan, 1972).

15. J. B. Losos, Lizards in an evolutionary tree: ecology and adaptive radiation of anoles (University of California Press, 2011).

16. A. Corl, A. R. Davis, S. R. Kuchta, T. Comendant, B. Sinervo, Alternative Mating Strategies and the Evolution of Sexual Size Dimorphism in the Side-Blotched Lizard, *Uta stansburiana*: A Population-Level Comparative Analysis. Evolution 64, 79–96 (2010).

17. B. D. Hollingsworth, “The Molecular Systematics of the Side-blotched Lizards (Iguania: Phrynosomatidae: Uta),” Loma Linda University. (1999).

18. D. E. Upton, R. W. Murphy, Phylogeny of the Side-Blotched Lizards (Phrynosomatidae: *Uta*) Based on mtDNA Sequences: Support for a Midpeninsular Seaway in Baja California. Molecular Phylogenetics and Evolution 8, 104–113 (1997).

19. K. M. Scoular, et al., Multiyear Home-Range Ecology of Common Side-blotched Lizards in Eastern Oregon with Additional Analysis of Geographic Variation in Home-range Size. Herpetological Monographs 25, 52–75 (2011).

20. B. S. Wilson, Latitudinal Variation in Activity Season Mortality Rates of the Lizard *Uta stansburiana*. Ecological Monographs 61, 393–414 (1991).

21. G. A. Dolby, S. E. K. Bennett, A. Lira-Noriega, B. T. Wilder, A. Munguía-Vega, Assessing the Geological and Climatic Forcing of Biodiversity and Evolution Surrounding the Gulf of California. *jsw* **57**, 391–455 (2015).

22. G. A. Dolby, et al., Integrating Earth–life systems: a geogenomic approach. Trends in Ecology & Evolution 37, 371–384 (2022).

23. K. Gardner, et al., Evidence for terrestrial origin of the pliocene San Regis beds, central Baja California peninsula, mexico in (2024).

24. R. Araya-Donoso, A. Biddy, A. Munguía-Vega, A. Lira-Noriega, G. A. Dolby, Habitat quality or quantity? Niche marginality across 21 plants and animals suggests differential responses between highland and lowland species to past climatic changes. Ecography 2024 (2024).

25. L. Cab-Sulub, S. T. Álvarez-Castañeda, Climatic dissimilarity associated with phylogenetic breaks. Journal of Mammalogy 102, 1592–1604 (2021).

26. R. Araya-Donoso, et al., Implications of barrier ephemerality in geogenomic research. Journal of Biogeography 49, 2050–2063 (2022).

27. A. D. Leaché, J. R. Oaks, C. Ofori-Boateng, M. K. Fujita, Comparative phylogeography of West African amphibians and reptiles. Evolution 74, 716–724 (2020).

28. R. S. Taylor, V. L. Friesen, The role of allochrony in speciation. Molecular Ecology 26, 3330–3342 (2017).

29. A. Klimova, et al., Metabarcoding reveals seasonal and spatial patterns of arthropod community assemblages in two contrasting habitats: Desert and oasis of the Baja California Peninsula, Mexico. Diversity and Distributions 29, 438–461 (2023).

30. H. A. Thomassen, A. H. Freedman, D. M. Brown, W. Buermann, D. K. Jacobs, Regional Differences in Seasonal Timing of Rainfall Discriminate between Genetically Distinct East African Giraffe Taxa. PLoS ONE 8, e77191 (2013).

31. C. E. Guarnizo, P. Montoya, I. Quintero, C. D. Cadena, Population divergence associated with spatial asynchrony in precipitation in Neotropical frogs. Journal of Biogeography 49, 2169–2180 (2022).

32. I. Quintero, S. González-Caro, P.-C. Zalamea, C. D. Cadena, Asynchrony of Seasons: Genetic Differentiation Associated with Geographic Variation in Climatic Seasonality and Reproductive Phenology. The American Naturalist 184, 352–363 (2014).

33. N. J. Mellor, et al., Divergence in Regulatory Regions and Gene Duplications May Underlie Chronobiological Adaptation in Desert Tortoises. Molecular Ecology 34, e17600 (2025).

34. W. S. Parker, E. R. Pianka, Comparative Ecology of Populations of the Lizard *Uta stansburiana*. Copeia 1975, 615 (1975).

35. P. A. Zani, S. J. Stein, Field and laboratory responses to drought by Common Side- blotched Lizards (Uta stansburiana). Journal of Arid Environments 154, 15–23 (2018).

36. R. Araya-Donoso, et al., Integrating genetics, physiology and morphology to study desert adaptation in a lizard species. Journal of Animal Ecology 91, 1148–1162 (2022).

37. J. L. Rocha, et al., North African fox genomes show signatures of repeated introgression and adaptation to life in deserts. Nat Ecol Evol 7, 1267–1286 (2023).

38. X. Peng, et al., Whole-genome sequencing reveals adaptations of hairy-footed jerboas (*Dipus*, Dipodidae) to diverse desert environments. BMC Biol 21 (2023).

39. H. Wu, et al., Camelid genomes reveal evolution and adaptation to desert environments. Nat Commun 5 (2014).

40. Cellular and Molecular Biology of Neuronal Dystonin (Elsevier, 2013).

41. M. Davenne, et al., Hoxa2 and Hoxb2 Control Dorsoventral Patterns of Neuronal Development in the Rostral Hindbrain. Neuron 22, 677–691 (1999).

42. S. J. James, S. Shpyleva, S. Melnyk, O. Pavliv, I. P. Pogribny, Complex epigenetic regulation of Engrailed-2 (EN-2) homeobox gene in the autism cerebellum. Transl Psychiatry 3, e232–e232 (2013).

43. M. Pagani, et al., Deletion of Autism Risk Gene Shank3 Disrupts Prefrontal Connectivity. J. Neurosci. 39, 5299–5310 (2019).

44. M. B. Passani, C. Ballerini, Histamine and neuroinflammation: insights from murine experimental autoimmune encephalomyelitis. Front. Syst. Neurosci. 6 (2012).

45. H. Indrischek, et al., Vision-related convergent gene losses reveal SERPINE3’s unknown role in the eye. eLife 11 (2022).

46. Y. Wada, J. Sugiyama, T. Okano, Y. Fukada, GRK1 and GRK7: Unique cellular distribution and widely different activities of opsin phosphorylation in the zebrafish rods and cones. Journal of Neurochemistry 98, 824–837 (2006).

47. D. R. Reed, et al., Polymorphisms in the Taste Receptor Gene (*Tas1r3*) Region Are Associated with Saccharin Preference in 30 Mouse Strains. J. Neurosci. 24, 938–946 (2004).

48. A. Tigano, J. P. Colella, M. D. MacManes, Comparative and population genomics approaches reveal the basis of adaptation to deserts in a small rodent. Molecular Ecology 29, 1300–1314 (2020).

49. J. M. Mwacharo, et al., Genomic footprints of dryland stress adaptation in Egyptian fat-tail sheep and their divergence from East African and western Asia cohorts. Sci Rep 7 (2017).

50. K. H. Cho, M. J. Kim, G. J. Jeon, H. Y. Chung, Association of genetic variants for FABP3 gene with back fat thickness and intramuscular fat content in pig. Mol Biol Rep 38, 2161–2166 (2011).

51. P. Aksoy, et al., Cytosolic 5′-nucleotidase III (NT5C3): gene sequence variation and functional genomics. Pharmacogenetics and Genomics 19, 567–576 (2009).

52. T. Desvignes, P. Pontarotti, C. Fauvel, J. Bobe, Nme protein family evolutionary history, a vertebrate perspective. BMC Evol Biol 9, 256 (2009).

53. Z. Jin, et al., Integrative multiomics evaluation reveals the importance of pseudouridine synthases in hepatocellular carcinoma. Front. Genet. 13, 944681 (2022).

54. J. Cheng, et al., Similar adaptative mechanism but divergent demographic history of four sympatric desert rodents in Eurasian inland. Commun Biol 6 (2023).

55. L. Xu, et al., Whole-genome resequencing provides insights into the diversity and adaptation to desert environment in Xinjiang Mongolian cattle. BMC Genomics 25 (2024).

56. L. G. McFerrin, W. R. Atchley, Evolution of the Max and Mlx Networks in Animals. Genome Biology and Evolution 3, 915–937 (2011).

57. B. Tepe, et al., Bi-allelic variants in INTS11 are associated with a complex neurological disorder. The American Journal of Human Genetics 110, 774–789 (2023).

58. P. Czerwińska, S. Mazurek, M. Wiznerowicz, The complexity of TRIM28 contribution to cancer. J Biomed Sci 24 (2017).

59. K. A. Wood, M. A. Eadsforth, W. G. Newman, R. T. O’Keefe, The Role of the U5 snRNP in Genetic Disorders and Cancer. Front. Genet. 12 (2021).

60. S. Yang, R. Jia, Z. Bian, SRSF5 functions as a novel oncogenic splicing factor and is upregulated by oncogene SRSF3 in oral squamous cell carcinoma. Biochimica et Biophysica Acta (BBA) - Molecular Cell Research 1865, 1161–1172 (2018).

61. N. Okamoto, et al., A novel genetic syndrome with *STARD9* mutation and abnormal spindle morphology. American J of Med Genetics Pt A 173, 2690–2696 (2017).

62. B. P. Duncker, I. N. Chesnokov, B. J. McConkey, The origin recognition complex protein family. Genome Biol 10, 214 (2009).

63. O. Markandona, et al., Single-nucleotide polymorphism rs 175080 in the MLH3 gene and its relation to male infertility. J Assist Reprod Genet 32, 1795–1799 (2015).

64. A. Dufner, S. Pownall, T. W. Mak, Caspase recruitment domain protein 6 is a microtubule-interacting protein that positively modulates NF-κB activation. Proc. Natl. Acad. Sci. U.S.A. 103, 988–993 (2006).

65. Z. Jiang, et al., RSL1D1 modulates cell senescence and proliferation via regulation of PPARγ mRNA stability. Life Sciences 307, 120848 (2022).

66. B. Ababaikeri, et al., Whole-genome sequencing of Tarim red deer (*Cervus elaphus yarkandensis*) reveals demographic history and adaptations to an arid-desert environment. Front Zool 17 (2020).

67. S. M. Baty, et al., Strong signatures of selection on genes underlying core reinforcement mechanisms in speciating desert tortoises. [Preprint] (2024). Available at: http://biorxiv.org/lookup/doi/10.1101/2024.06.06.597788 [Accessed 1 October 2024].

68. C. E. Chapin, Interplay of oceanographic and paleoclimate events with tectonism during middle to late Miocene sedimentation across the southwestern USA. Geosphere 4, 976 (2008).

69. M. H. Darin, et al., Revised age Depositional age of the Boleo Formation and marine flooding in the central Gulf of California rift, Santa Rosalía basin, Baja California Sur, México. Geol. Soc. Am. Spec. Pap. (2024).

70. T. Bhattacharya, et al., Expansion and Intensification of the North American Monsoon During the Pliocene. AGU Advances 3, e2022AV000757 (2022).

71. M. M. Sparks, C. E. Schraidt, X. Yin, L. W. Seeb, M. R. Christie, Rapid genetic adaptation to a novel ecosystem despite a large founder event. Molecular Ecology (2023). 10.1111/mec.17121.

72. F. Termignoni-Garcia, J. J. Kirchman, J. Clark, S. V. Edwards, Comparative Population Genomics of Cryptic Speciation and Adaptive Divergence in Bicknell’s and Gray-Cheeked Thrushes (Aves: *Catharus bicknelli* and *Catharus minimus*). Genome Biology and Evolution 14 (2022).

73. M. Goller, F. Goller, S. S. French, A heterogeneous thermal environment enables remarkable behavioral thermoregulation in *Uta stansburiana*. Ecology and Evolution 4, 3319–3329 (2014).

74. S. Waldschmidt, C. R. Tracy, Interactions between a Lizard and Its Thermal Environment: Implications for Sprint Performance and Space Utilization in the Lizard *Uta stansburiana*. Ecology 64, 476–484 (1983).

75. S. C. Campbell-Staton, A. Bare, J. B. Losos, S. V. Edwards, Z. A. Cheviron, Physiological and regulatory underpinnings of geographic variation in reptilian cold tolerance across a latitudinal cline. Molecular Ecology 27, 2243–2255 (2018).

76. S. Fellows, et al., Chromosome-length genome assembly of Uta stansburiana and gene expression data reveal fast pace-of-life comes with environmental stability. [Preprint] (2025). Available at: http://biorxiv.org/lookup/doi/10.1101/2025.05.28.656178 [Accessed 2 July 2025].

77. R Core Team, R: A language and environment for statistical computing. (2022). Deposited 2022.

78. K. Mokany, C. Ware, S. N. C. Woolley, S. Ferrier, M. C. Fitzpatrick, A working guide to harnessing generalized dissimilarity modelling for biodiversity analysis and conservation assessment. Global Ecol Biogeogr 31, 802–821 (2022).

79. D. G. Moore, M. Morales, A. R. Biddy, S. I. Walker, G. A. Dolby, The information signature of diverging lineages. [Preprint] (2021). Available at: http://biorxiv.org/lookup/doi/10.1101/2021.08.30.458276 [Accessed 9 April 2025].

## References

1. S. Chen, Ultrafast one-pass FASTQ data preprocessing, quality control, and deduplication using fastp. iMeta 2 (2023).

2. S. Fellows, et al., Chromosome-length genome assembly of Uta stansburiana and gene expression data reveal fast pace-of-life comes with environmental stability. [Preprint] (2025). Available at: http://biorxiv.org/lookup/doi/10.1101/2025.05.28.656178 [Accessed 2 July 2025].

3. H. Li, R. Durbin, Fast and accurate short read alignment with Burrows–Wheeler transform. Bioinformatics 25, 1754–1760 (2009).

4. A. McKenna, et al., The Genome Analysis Toolkit: A MapReduce framework for analyzing next-generation DNA sequencing data. Genome Res. 20, 1297–1303 (2010).

5. G. A. Van Der Auwera, et al., From FastQ Data to High-Confidence Variant Calls: The Genome Analysis Toolkit Best Practices Pipeline. CP in Bioinformatics 43 (2013).

6. P. Danecek, et al., The variant call format and VCFtools. Bioinformatics 27, 2156– 2158 (2011).

7. S. Purcell, et al., PLINK: A Tool Set for Whole-Genome Association and Population- Based Linkage Analyses. The American Journal of Human Genetics 81, 559–575 (2007).

8. R Core Team, R: A language and environment for statistical computing. (2022). Deposited 2022.

9. D. H. Alexander, J. Novembre, K. Lange, Fast model-based estimation of ancestry in unrelated individuals. Genome Res. 19, 1655–1664 (2009).

10. D. Petkova, J. Novembre, M. Stephens, Visualizing spatial population structure with estimated effective migration surfaces. Nat Genet 48, 94–100 (2016).

11. C. J. Versoza, et al., The recombination landscapes of spiny lizards (genus *Sceloporus*). G3 Genes|Genomes|Genetics 12, jkab402 (2022).

12. S. Sánchez, vcf2fasta.

13. D. Darriba, G. L. Taboada, R. Doallo, D. Posada, jModelTest 2: more models, new heuristics and parallel computing. Nat Methods 9, 772–772 (2012).

14. M. A. Suchard, et al., Bayesian phylogenetic and phylodynamic data integration using BEAST 1.10. Virus Evolution 4 (2018).

15. S. Kumar, G. Stecher, M. Suleski, S. B. Hedges, TimeTree: A Resource for Timelines, Timetrees, and Divergence Times. Molecular Biology and Evolution 34, 1812–1819 (2017).

16. J. Josse, F. Husson, missMDA: A Package for Handling Missing Values in Multivariate Data Analysis. J. Stat. Soft. 70 (2016).

17. M. Rizzo, G. Szekely, energy: E-Statistics: Multivariate Inference via the Energy of Data. 10.32614/CRAN.package.energy. Deposited 6 February 2004.

18. J. Oksanen, et al., vegan: Community Ecology Package. 10.32614/CRAN.package.vegan. Deposited 6 September 2001.

19. GBIF.org, Occurrence Download. The Global Biodiversity Information Facility. 10.15468/DL.HGEJX4. Deposited 2024.

20. S. E. Fick, R. J. Hijmans, WorldClim 2: new 1-km spatial resolution climate surfaces for global land areas. Intl Journal of Climatology 37, 4302–4315 (2017).

21. QGIS.org, QGIS Geographic Information System. (2020). Deposited 2020.

22. M. E. Cobos, L. Osorio-Olvera, J. Soberón, A. T. Peterson, N. Barve, ellipsenm: ecological niche’s characterizations using ellipsoids - R package. (2020). Deposited 2020.

23. J. Van Etten, *R* Package gdistance: Distances and Routes on Geographical Grids. J. Stat. Soft. 76 (2017).

24. K. Mokany, C. Ware, S. N. C. Woolley, S. Ferrier, M. C. Fitzpatrick, A working guide to harnessing generalized dissimilarity modelling for biodiversity analysis and conservation assessment. Global Ecol Biogeogr 31, 802–821 (2022).

25. M. Fitzpatrick, K. Mokany, G. Manion, D. Nieto-Lugilde, S. Ferrier, gdm: Generalized Dissimilarity Modeling. 10.32614/CRAN.package.gdm. Deposited 27 March 2015.

105. D. G. Moore, M. Morales, A. R. Biddy, S. I. Walker, G. A. Dolby, The information signature of diverging lineages. [Preprint] (2021). Available at: http://biorxiv.org/lookup/doi/10.1101/2021.08.30.458276 [Accessed 9 April 2025].

27. P. L. Williams, R. D. Beer, Nonnegative Decomposition of Multivariate Information. [Preprint] (2010). Available at: http://arxiv.org/abs/1004.2515 [Accessed 18 January 2023].

28. M. Harder, C. Salge, D. Polani, A Bivariate Measure of Redundant Information. *Phys*. Rev. E 87, 012130 (2013).

29. J. Kay, J. Schulz, W. Phillips, A Comparison of Partial Information Decompositions Using Data from Real and Simulated Layer 5b Pyramidal Cells. Entropy 24, 1021 (2022).

30. J. D. Scargle, J. P. Norris, B. Jackson, J. Chiang, STUDIES IN ASTRONOMICAL TIME SERIES ANALYSIS. VI. BAYESIAN BLOCK REPRESENTATIONS. ApJ 764, 167 (2013).

31. M. Kochenderfer, 2025, Discretizers. https://github.com/sisl/Discretizers.jl.

32. D. Moore, 2021, Imogen. https://github.com/elife-asu/Imogen.jl.

33. J. Terhorst, J. A. Kamm, Y. S. Song, Robust and scalable inference of population history from hundreds of unphased whole genomes. Nat Genet 49, 303–309 (2017).

34. X. Liu, Y.-X. Fu, Stairway Plot 2: demographic history inference with folded SNP frequency spectra. Genome Biol 21 (2020).

35. Y. Bourgeois, R. P. Ruggiero, J. D. Manthey, S. Boissinot, Recent Secondary Contacts, Linked Selection, and Variable Recombination Rates Shape Genomic Diversity in the Model Species *Anolis carolinensis*. Genome Biology and Evolution 11, 2009–2022 (2019).

36. F. B. Turner, G. A. Hoddenbach, P. A. Medica, J. R. Lannom, The Demography of the Lizard, *Uta stansburiana* Baird and Girard, in Southern Nevada. The Journal of Animal Ecology 39, 505 (1970).

37. V. Obenchain, et al., VariantAnnotation : a Bioconductor package for exploration and annotation of genetic variants. Bioinformatics 30, 2076–2078 (2014).

38. R. Andersson, A. Sandelin, Determinants of enhancer and promoter activities of regulatory elements. Nat Rev Genet 21, 71–87 (2020).

39. D. J. Wilson, et al., GenomegaMap: Within-Species Genome-Wide dN/dS Estimation from over 10,000 Genomes. Molecular Biology and Evolution 37, 2450–2460 (2020).

40. L. Kolberg, et al., g:Profiler—interoperable web service for functional enrichment analysis and gene identifier mapping (2023 update). Nucleic Acids Research 51, W207–W212 (2023).

41. D. Szklarczyk, et al., STRING v11: protein–protein association networks with increased coverage, supporting functional discovery in genome-wide experimental datasets. Nucleic Acids Research 47, D607–D613 (2019).

42. G. Stelzer, et al., The GeneCards Suite: From Gene Data Mining to Disease Genome Sequence Analyses. CP in Bioinformatics 54 (2016).

43. The UniProt Consortium, et al., UniProt: the Universal Protein Knowledgebase in 2025. Nucleic Acids Research 53, D609–D617 (2025).

