## Supplemental Information for "Behavioral and phenotypic constraint belie deep genomic divergence and seasonal adaptation in a widespread desert lizard"

18. J. Oksanen, *et al.*, vegan: Community Ecology Package. https://doi.org/10.32614/CRAN.package.vegan. Deposited 6 September 2001.

19. GBIF.org, Occurrence Download. The Global Biodiversity Information Facility. https://doi.org/10.15468/DL.HGEJX4. Deposited 2024.

20. S. E. Fick, R. J. Hijmans, WorldClim 2: new 1-km spatial resolution climate surfaces for global land areas. *Intl Journal of Climatology* **37**, 4302–4315 (2017).

21. QGIS.org, QGIS Geographic Information System. (2020). Deposited 2020.

22. M. E. Cobos, L. Osorio-Olvera, J. Soberón, A. T. Peterson, N. Barve, ellipsenm: ecological niche's characterizations using ellipsoids - R package. (2020). Deposited 2020.

23. J. Van Etten, *R* Package gdistance: Distances and Routes on Geographical Grids. *J. Stat. Soft.* **76** (2017).

24. K. Mokany, C. Ware, S. N. C. Woolley, S. Ferrier, M. C. Fitzpatrick, A working guide to harnessing generalized dissimilarity modelling for biodiversity analysis and conservation assessment. *Global Ecol Biogeogr* **31**, 802–821 (2022).

25. M. Fitzpatrick, K. Mokany, G. Manion, D. Nieto-Lugilde, S. Ferrier, gdm: Generalized Dissimilarity Modeling. https://doi.org/10.32614/CRAN.package.gdm. Deposited 27 March 2015.

26. D. G. Moore, M. Morales, A. R. Biddy, S. I. Walker, G. A. Dolby, The information signature of diverging lineages. [Preprint] (2021). Available at: http://biorxiv.org/lookup/doi/10.1101/2021.08.30.458276 [Accessed 9 April 2025].

### Supplementary Figures

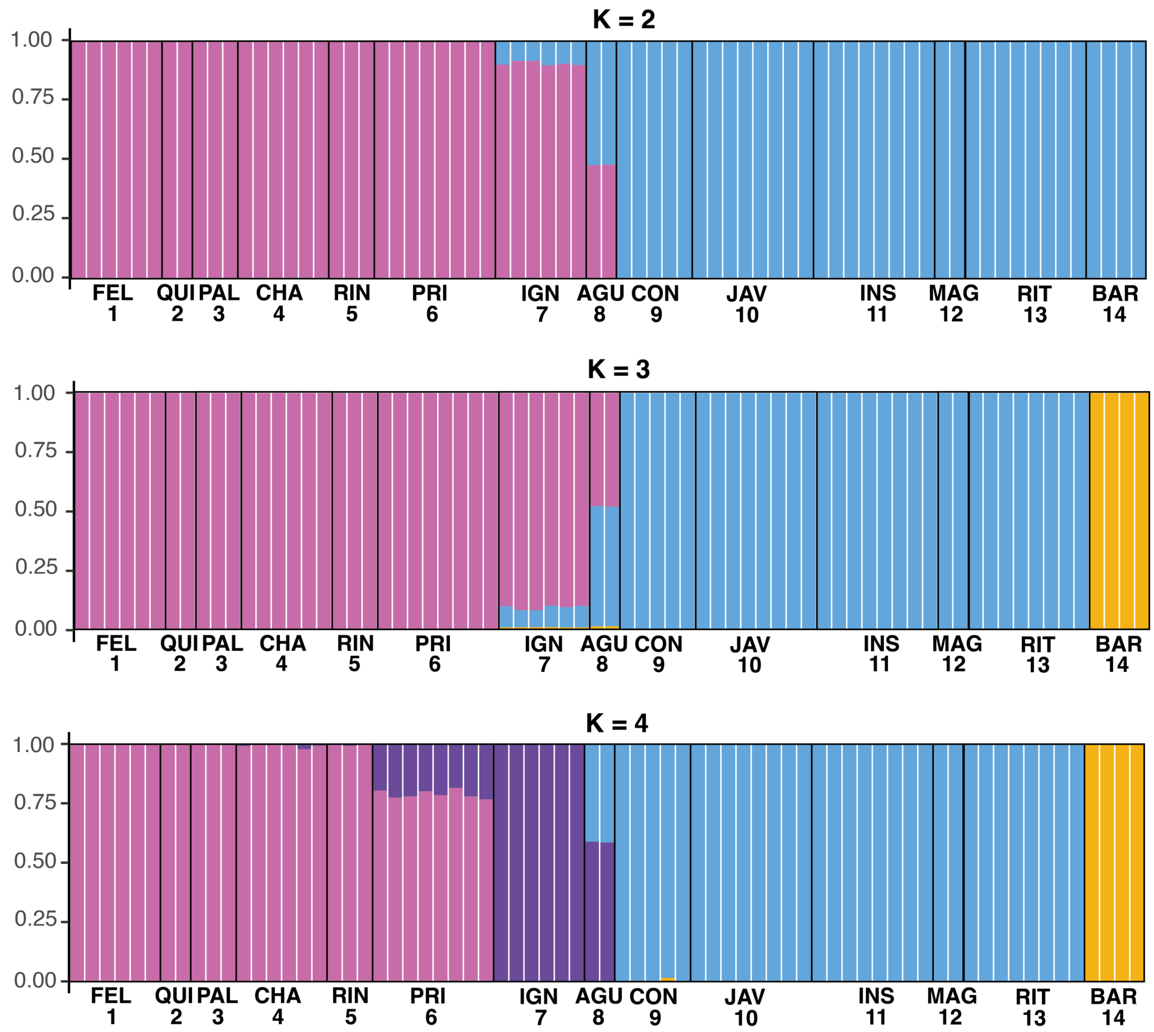


**Figure S1.** Structure plots for K = 2 to K = 4 for *Uta stansburiana* populations on the Baja California peninsula.

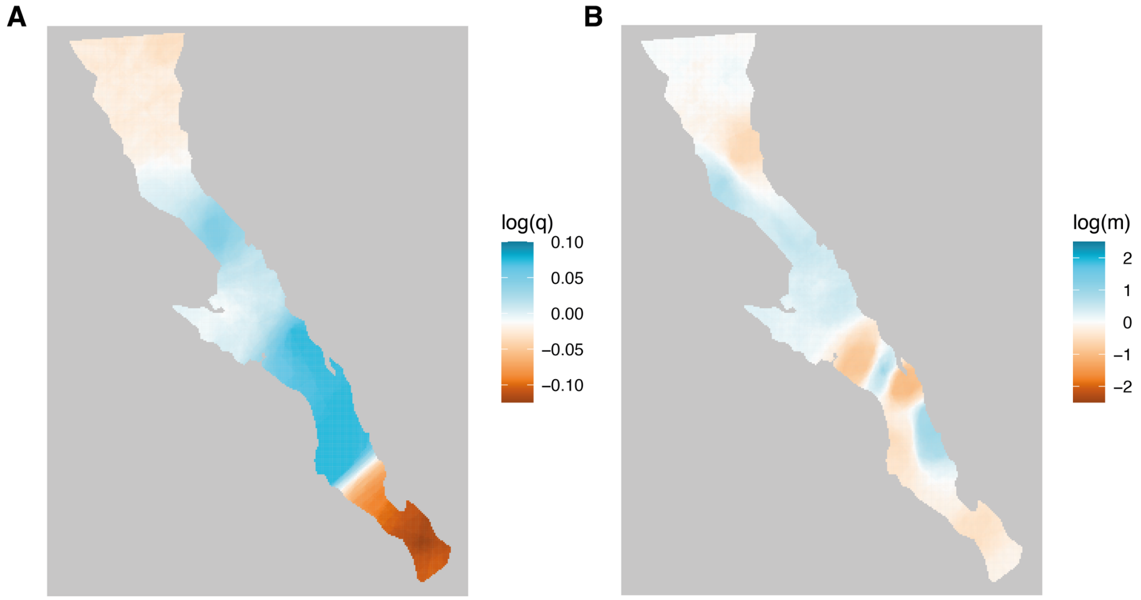


**Figure S2.** Effective genetic diversity (**A**) and migration (**B**) surfaces for Uta stansburiana on the Baja California peninsula.

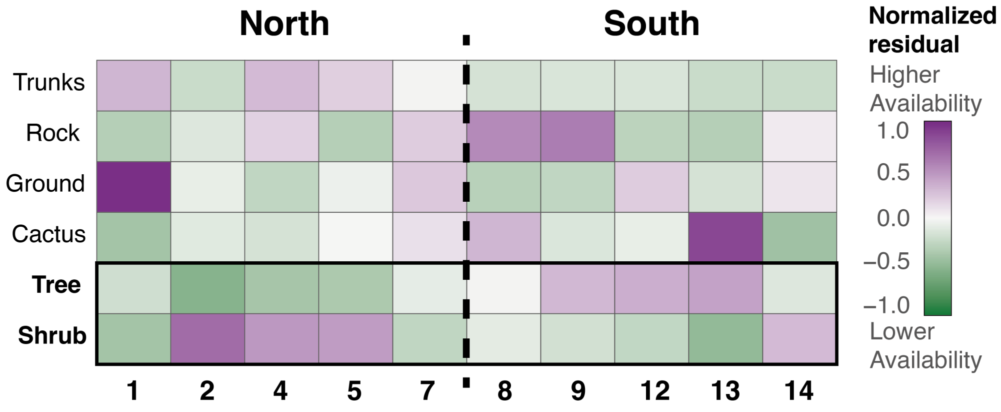


**Figure S3.** Habitat availability for ten sampling locations of *Uta stansburiana* on the Baja California peninsula. In general, northern sites have higher shrub cover whereas southern sites have higher tree cover.

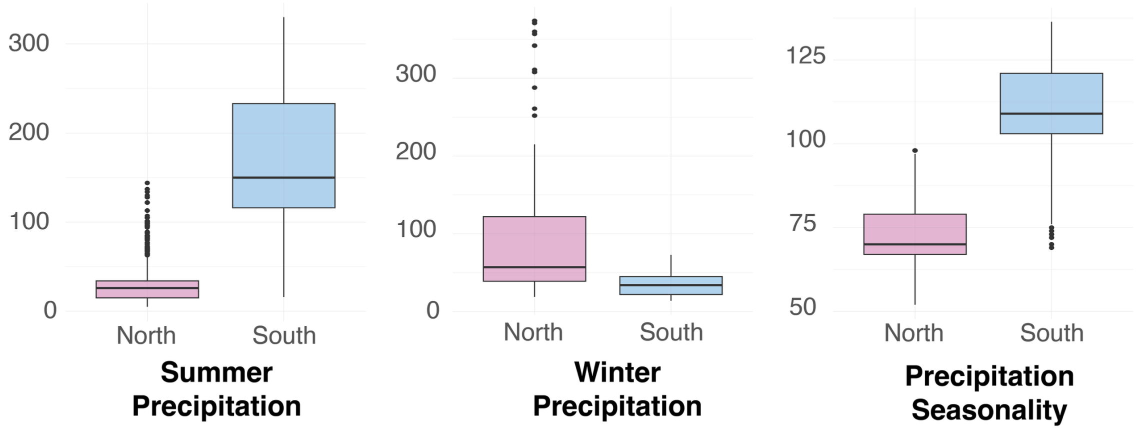


**Figure S4.** Variation in precipitation variables (mm) for the northern and southern locations on the Baja California peninsula.

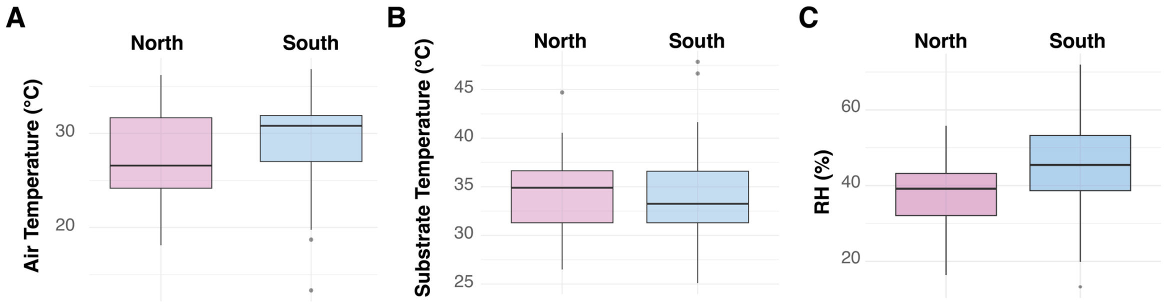


**Figure S5. A.** Air temperature, **B.** Substrate temperature, and **C.** Ambient relative humidity for *Uta stansburiana* from the northern and southern genetic groups on the Baja California peninsula.

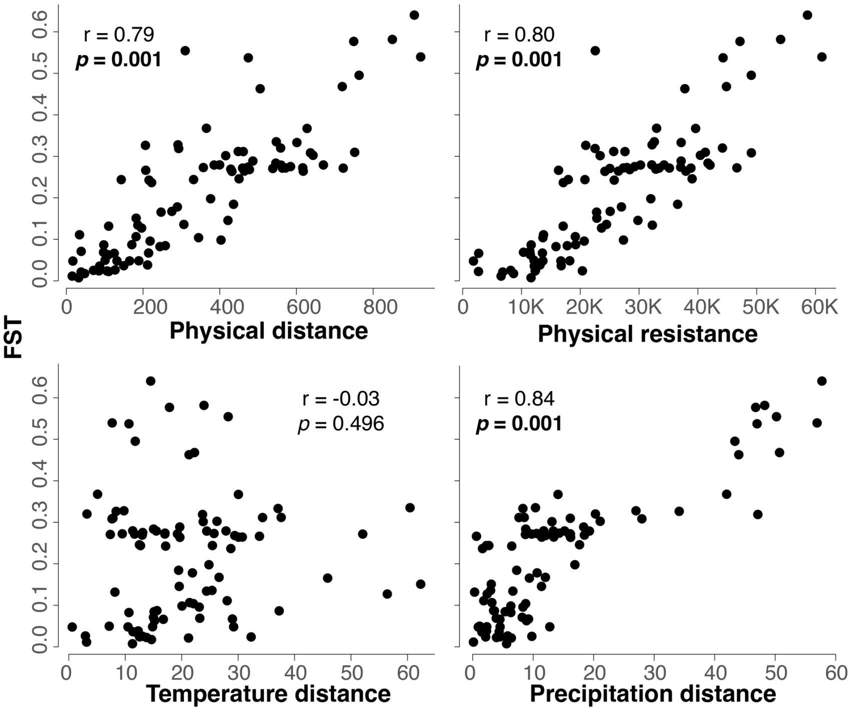


**Figure S6.** Mantel tests for isolation by distance, isolation by topographic resistance and isolation by temperature and precipitation distance.

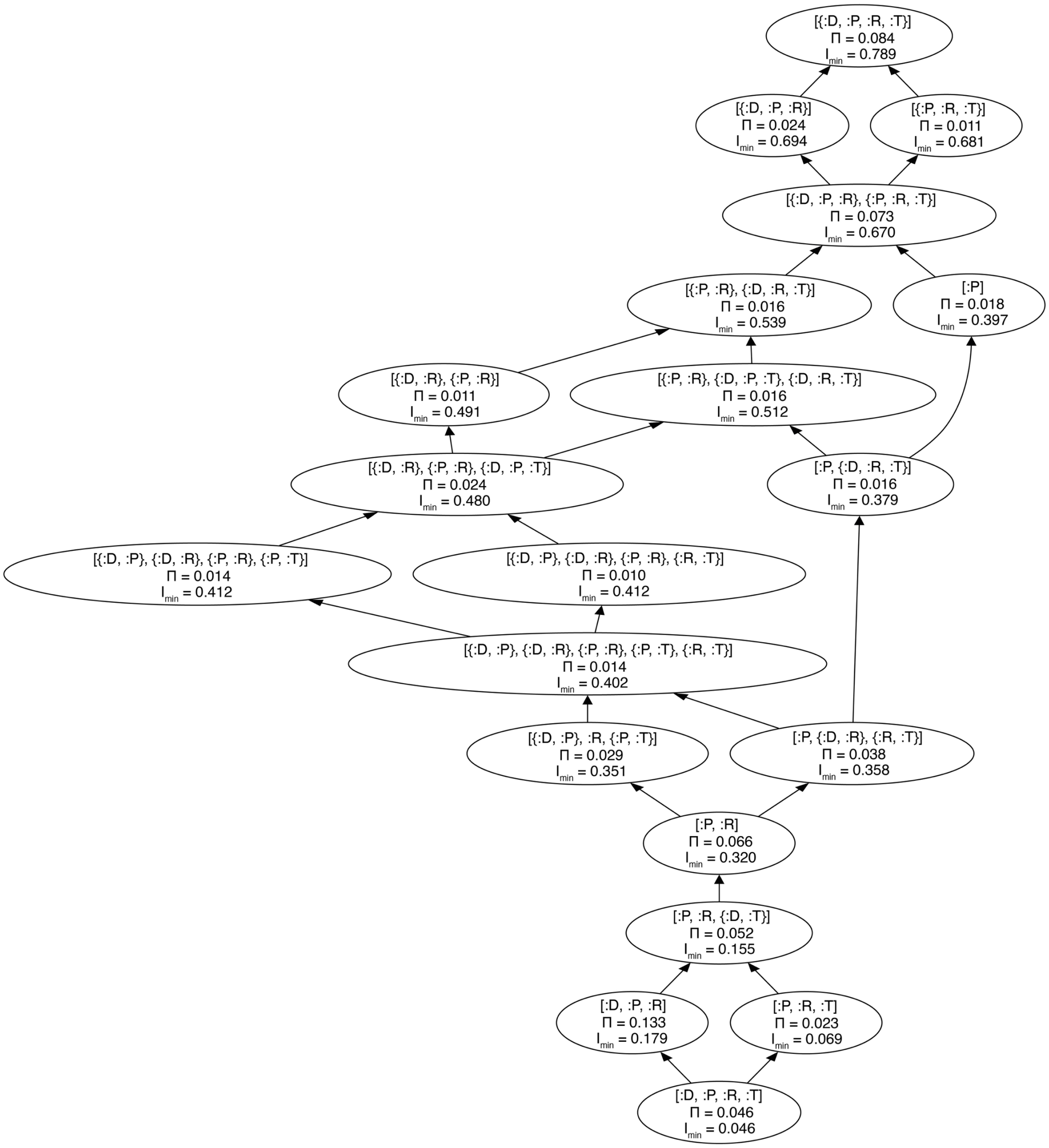


**Figure S7.** Partial information decomposition lattice for between-population $F_{ST}$ for Uta stansburiana. Each node in the lattice represents the information contribution of a combination of environmental distance predictors (D: distance, R: resistance, T: temperature, P: precipitation). In the lattice, [:X, :Y] represents redundant information between variables X and Y, whereas [{:X, :Y}] represents synergistic information between X and Y.

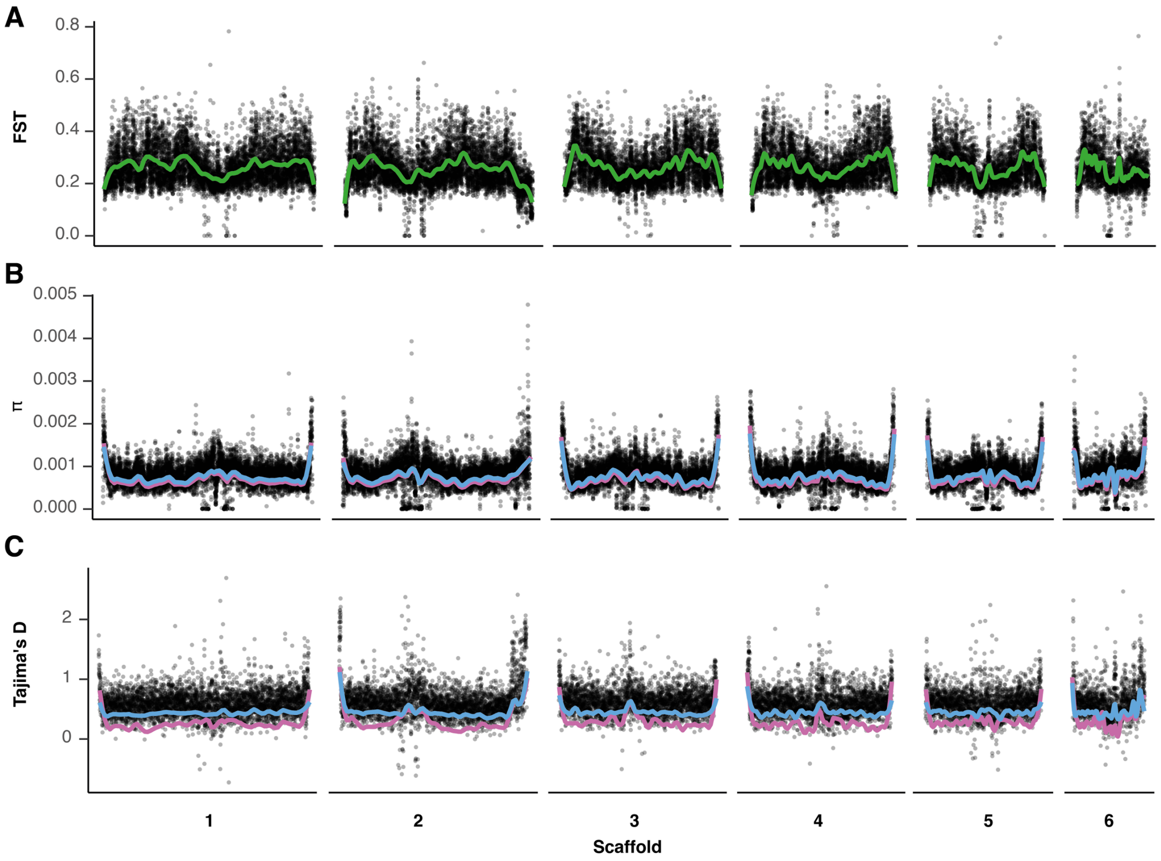


**Figure S8.** Population genetic statistics calculated along the genome for both genetic groups of *U. stansburiana*. Plot shows values for scaffolds 1-6 corresponding to the macrochromosomes. The northern genetic group is in pink, and the southern in blue. **A.** $F_{ST}$. **B.** $\pi$. **C.** Tajima's D.

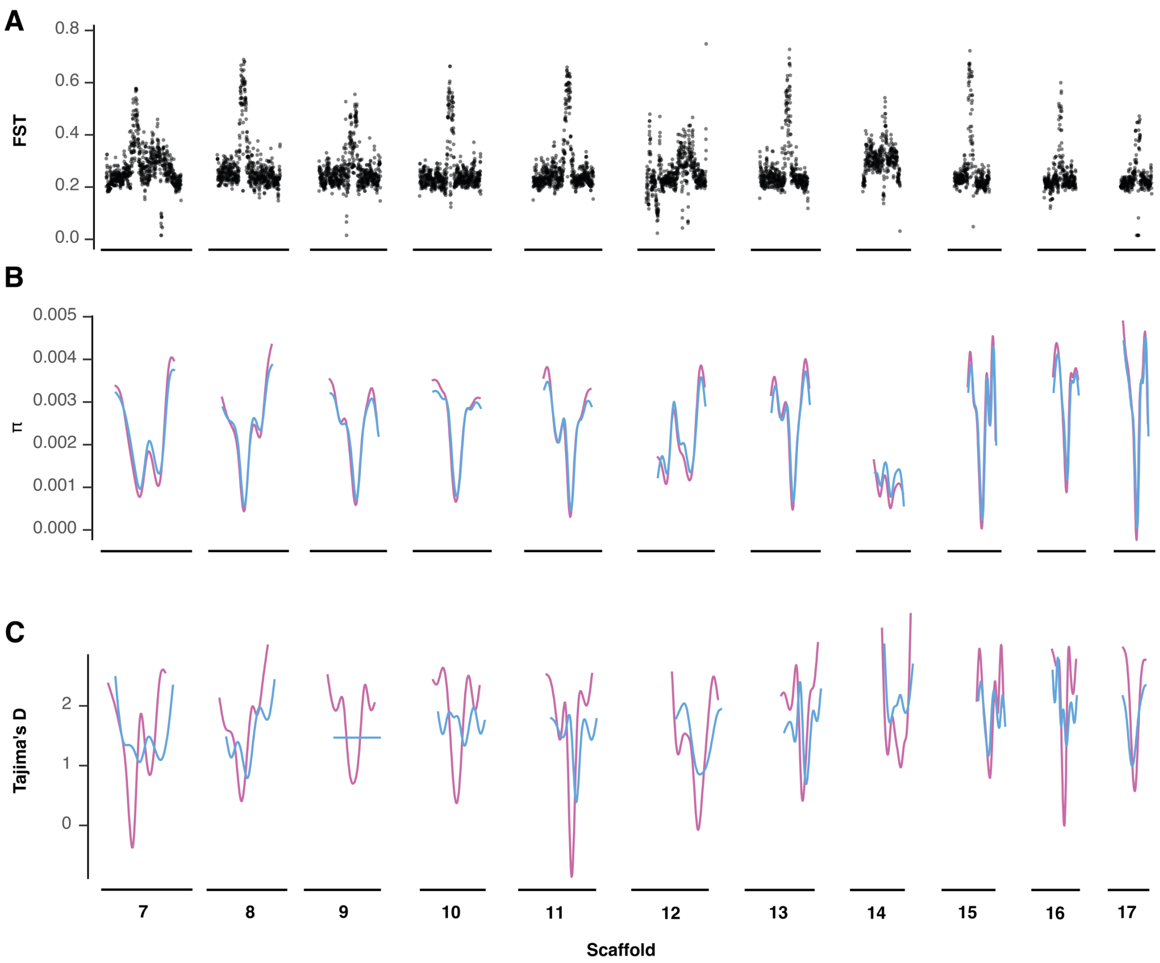


333

334 **Figure S9.** $F_{ST}$, Nucleotide diversity and Tajima's D for the micro-chromosomes of *Uta*
335 *stansburiana* populations on the Baja California peninsula. The northern genetic group is in
336 pink, and the southern in blue.

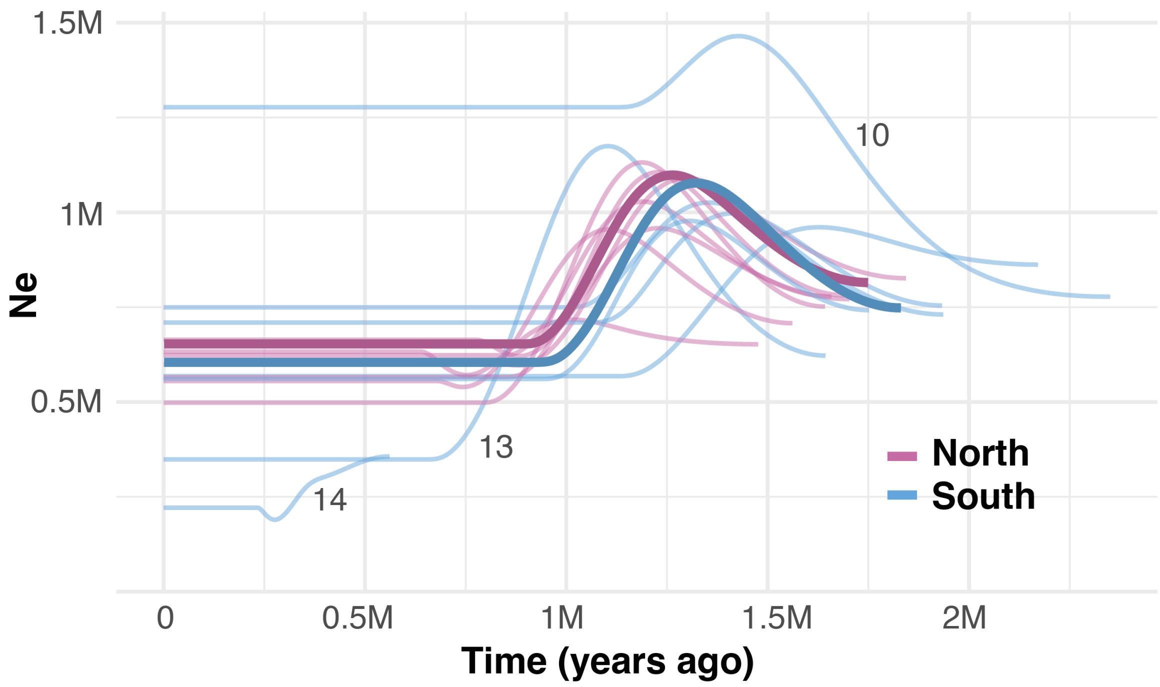


**Figure S10.** Past demography reconstruction with SMC++ for each population (light colors) and genetic group (darker colors) of U. stansburiana on the Baja California peninsula.

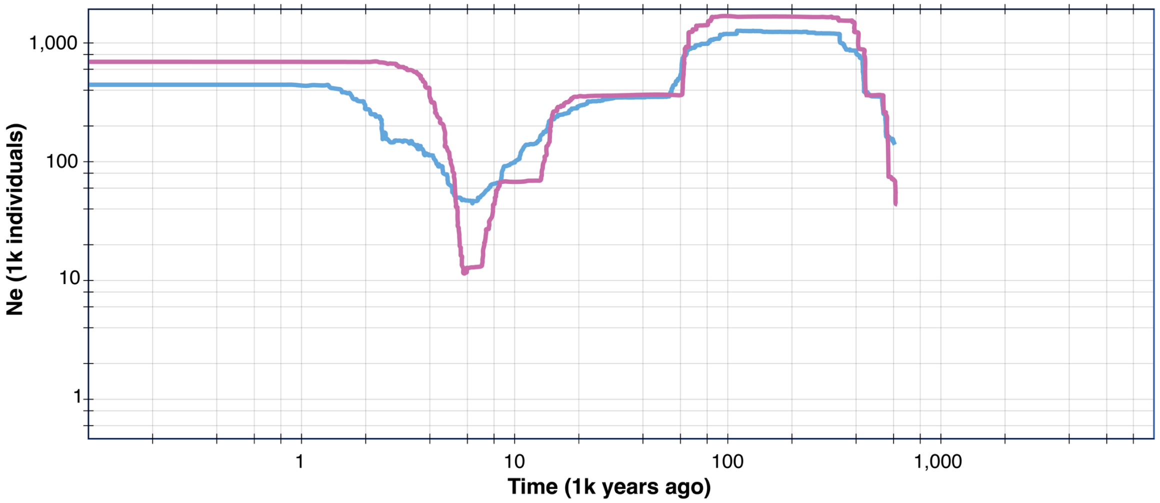


**Figure S11.** Demographic analysis performed with Stairwayplot on both genetic groups of *Uta stansburiana* on the Baja California peninsula.

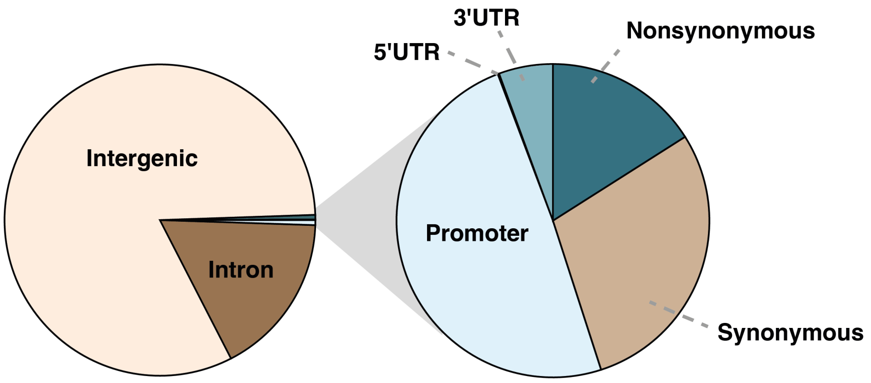


**Figure S12.** Functional classification of variants located within the highest 1% $F_{ST}$ regions between northern and southern genetic groups of *Uta stansburiana*.

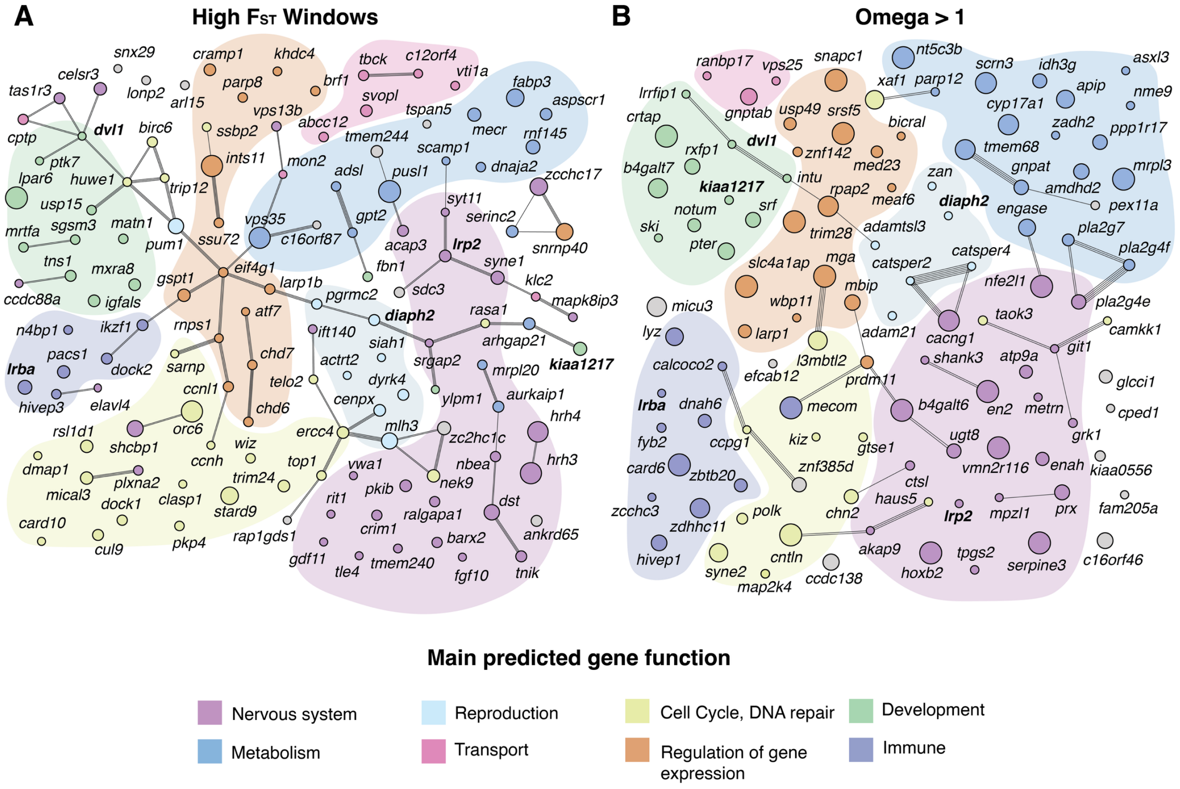


**Figure S13. A.** String network analysis for genes from the high FST regions with variants located in their promoter region, or non-synonymous variants in their coding regions. **B.** String network analysis for genes with ω > 1, and one allele fixed in one of the genetic groups. Lines represent known interactions between genes, and colors indicate the potential function of each gene. Circle size is proportional to the number of SNPs located in each coding sequence or promoter region in **A**, and the ω value in **B**. Genes overlapping between methodologies are highlighted in bold.

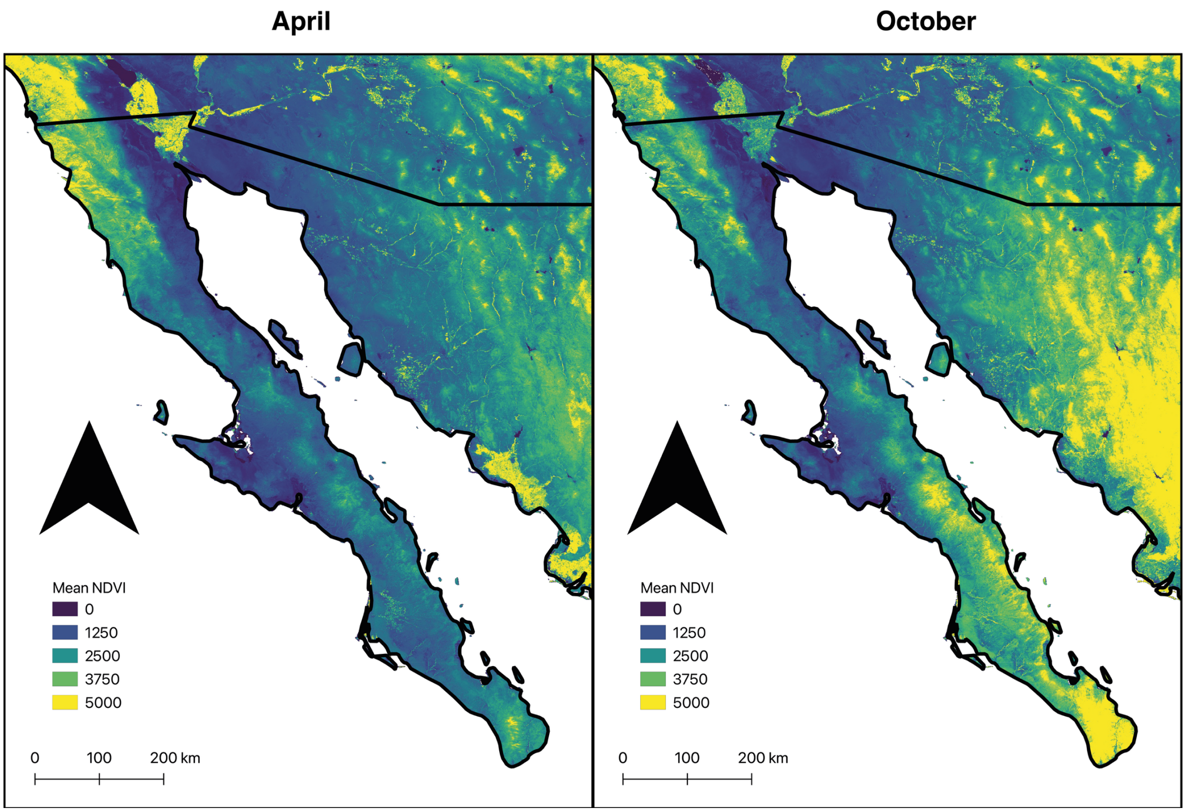


**Figure S14.** Seasonal variation in vegetation green-up time along the Baja California peninsula showed as the average NDVI values between 1993-2023 for April and October.

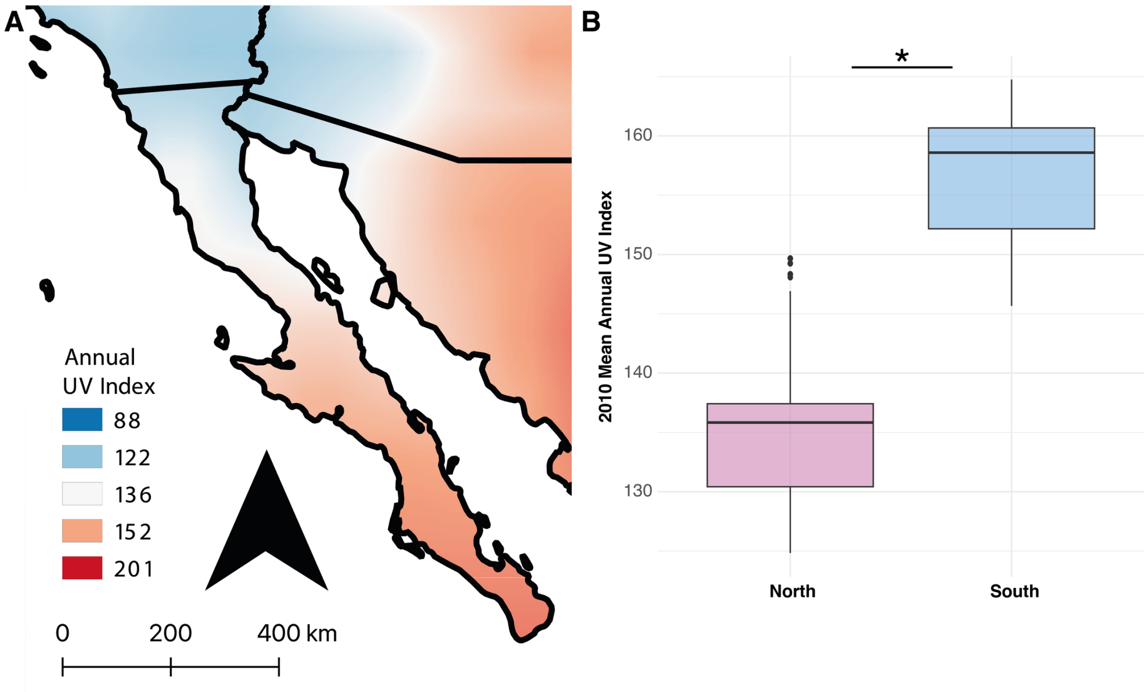


**Figure S15. A.** Variation of UV radiation across the Baja California peninsula. **B.** The northern and southern regions have significantly ($p < 0.001$) different annual UV indices.
